# Glaucoma-associated Optineurin mutations increase transmitophagy in a vertebrate optic nerve

**DOI:** 10.1101/2023.05.26.542507

**Authors:** Yaeram Jeong, Chung-ha O. Davis, Aaron M. Muscarella, Viraj Deshpande, Keun-Young Kim, Mark H. Ellisman, Nicholas Marsh-Armstrong

## Abstract

We previously described a process referred to as transmitophagy where mitochondria shed by retinal ganglion cell (RGC) axons are transferred to and degraded by surrounding astrocytes in the optic nerve head of mice. Since the mitophagy receptor Optineurin (OPTN) is one of few large-effect glaucoma genes and axonal damage occurs at the optic nerve head in glaucoma, here we explored whether OPTN mutations perturb transmitophagy. Live-imaging of *Xenopus laevis* optic nerves revealed that diverse human mutant but not wildtype OPTN increase stationary mitochondria and mitophagy machinery and their colocalization within, and in the case of the glaucoma-associated OPTN mutations also outside of, RGC axons. These extra-axonal mitochondria are degraded by astrocytes. Our studies support the view that in RGC axons under baseline conditions there are low levels of mitophagy, but that glaucoma-associated perturbations in OPTN result in increased axonal mitophagy involving the shedding and astrocytic degradation of the mitochondria.

**Graphical Abstract:** 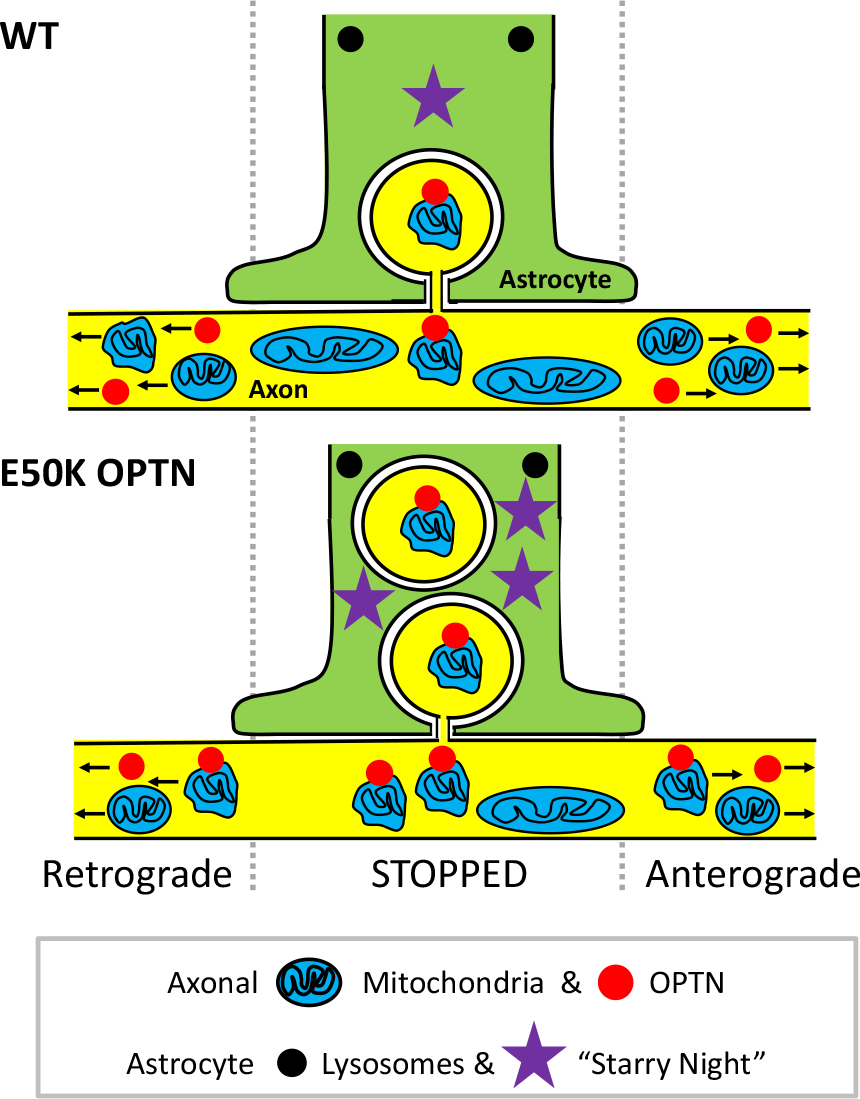

## Introduction

Mitochondria carry out diverse functions including being the major source of energy production (Attwell and Laughlin, 2001), and thus the proper quality control of mitochondria is essential for viability (Chen and Chan, 2009; Schon and Przedborski, 2011; Rugarli and Langer, 2012). Mitophagy entails either the constitutive or induced degradation of damaged or unnecessary mitochondria using highly conserved autophagic pathways that are also used to degrade other organelles, aggregates and pathogens (Ashrafi and Schwarz, 2013; Ding and Yin, 2012; Youle and Narendra, 2011). Mitochondria are often targeted for degradation by a process regulated by a sequence of phosphorylation and ubiquitination steps regulated by two genes that have been linked to Parkinson’s Disease, PTEN-induced putative kinase 1 (PINK1) and Parkin (Kitada et al., 1998;

Narendra et al., 2008). Optineurin (OPTN) is one of the principal mitophagy receptors that recognizes these ubiquitinated mitochondria (Wong and Holzbaur, 2014a; Heo et al., 2015; Lazarou et al., 2015), others being Nuclear dot protein 52 kDa (NDP52), neighbor of BRCA1 gene 1 (NBR1) and Sequestosome-1 (SQSTM1 or p62) (Rogov et al., 2014). Once OPTN is recruited to the ubiquitinated mitochondria, these mitochondria can then associate with microtubule-associated protein 1A/1B-light chain 3 (LC3) on nascent autophagosomes through the LC3 interacting region of OPTN, leading to full engulfment of the tagged organelles inside the autophagosomes, followed by lysosomal fusion that enables the subsequent degradation of the contained mitochondria (Stolz et al., 2014).

Although mitochondria are primarily produced and degraded in or near the cell soma (Burke et al., 1988; Saxton and Hollenbeck, 2012), many neuronal mitochondria also reside in distal processes far from the cell body (Nafstad and Blackstad, 1966), a situation that is the most extreme in the axons of long projection neurons such as RGCs (Yu et al., 2013). To date, most studies have focused on mitochondria quality control mechanisms in the soma of cells, and much less is known about mitochondria quality control in axons. Since effective clearance of damaged organelles in long axons is essential for the maintenance of the integrity of neurons, it has been a long-standing question whether adequate mitochondria quality control, more specifically efficient removal of dysfunctional mitochondria, can occur at these distal processes (Lu, 2014). There are studies that showed that distal axonal damaged mitochondria, or alternatively damaged parts of mitochondria that undergo fission from the rest of the mitochondria, are transported retrogradely for either lysosomal degradation, or alternatively fusion with healthy mitochondria in or near the soma (Hollenbeck, 1993; Miller and Sheetz, 2004; Maday et al., 2012; Maday and Holzbaur, 2014; Cheng et al., 2015; Zheng et al., 2019; Evans and Holzbaur, 2020). Consistent with these studies, there is evidence that axonal autophagy initiates preferentially at the distal tips or presynaptic sites of axons and terminates the clearance process at the soma (Yue, 2007; Wang et al., 2015; Soukup et al., 2016; Stavoe et al., 2016), suggesting that ultimate degradation mostly occurs in or near the soma, and thus leaving it an open question whether completion of axonal autophagy or mitophagy can ever occur locally within axons. Other studies have demonstrated that axonal mitochondria in mid- or distal- axons tend to be older and more susceptible to damage compared to those residing in the proximal axons or cell body (Lehmann et al., 2011; Ferree et al., 2013), and the vulnerable or stressed axonal mitochondria have diminished mitochondrial motility and thus fail to return to the soma for degradation (Wang et al., 2011; Lovas and Wang, 2013; Ashrafi et al., 2014; Hsieh et al., 2016; Cheng and Sheng, 2021), implying that not only initiating but also completing mitophagy locally in the axon is needed to remove the arrested old or damaged mitochondria effectively from long axons. Indeed, *Pink1* mRNA can be transported to distal regions of neurons and translated locally, providing a constant supply of fresh PINK1 protein for axonal mitochondria, and that this mechanism enables mitochondria to be degraded locally in the distal regions of axons (Harbauer et al., 2022).

We previously reported that in the optic nerve head of mice an alternative mechanism exists to degrade axonal mitochondria, whereby outpocketings of axons containing mitochondria are pinched off from axons and are degraded by the local astrocytes (Davis et al., 2014). Although they occurred elsewhere in the central nervous system, including in the cerebral cortex, these sheddings were most numerous in the optic nerve head, which is both a highly specialized anatomical structure where axons are constantly subject to biomechanical stress and is also the site of axon damage in glaucoma (Burgoyne et al., 2005; Sigal et al., 2005). More recently, similar sheddings of mitochondria have been observed in both Parkinson’s and Alzheimer’s Disease rodent animal models (Morales et al., 2020; Lampinen et al., 2022) as well as in cone photoreceptors (Hutto et al., 2023). What we initially described as axonal protrusions and evulsions likely represent a variant of what others have more recently described as exophers (Melentijevic et al., 2017; Nicolás-Ávila et al., 2020), large cellular outpocketing and sheddings which form in response to varied stressors and which contain various contents, including mitochondria.

OPTN is one of the very few genes that have a large effect on glaucoma (Rezaie et al., 2002; Meng et al., 2012), and the mutations in OPTN that promote glaucoma but not those that promote Amyotrophic Lateral Sclerosis (ALS) appear to promote more or possibly dysregulated mitophagy (Wong and Holzbaur, 2014a; Sirohi et al., 2015; Shim et al., 2016). Since mutations in mitophagy machinery result in damage to axons in the optic nerve head (Minegishi et al., 2016), and this is the location with the highest amounts of transmitophagy (Davis et al., 2014), the current study sought to determine whether mutations in OPTN associated with glaucoma might lead to alterations in transmitophagy. Since much of what has been learned about OPTN in mitophagy derives from live-imaging studies of mitochondria and mitophagic machinery in cultured cells (Lazarou et al., 2015; Moore and Holzbaur, 2016; Shim et al., 2016; Evans and Holzbaur, 2020), we took a similar live-imaging approach, though in this case in live animals where the anatomical relationship between axons and their cellular neighbors is maintained, as these cell-cell interactions are likely necessary to reproduce the relevant cell biology of transmitophagy. These studies were carried out in young *Xenopus laevis* tadpoles, as this vertebrate model system has two properties optimal for the proposed studies, an optic nerve that can be readily live-imaged and a highly efficient transgenesis suitable for the interrogation of gene variants.

## Results

### Live-imaging of mitochondria in RGC axons shows that under basal conditions approximately half of axonal mitochondria are stopped

To live-image RGC mitochondria within axons, we first intravitreally injected Mitotracker Deep Red into Nieuwkoop and Faber ((Nieuwkoop and Faber, 1994)) stage 48 (10 days post-fertilization) *X. laevis* tadpoles, animals whose RGC axons were readily visible due to expression of an RGC-specific *Tg(Isl2b:GFP)* transgene (Fig. 1A). After allowing the dye to spread down the axons, the optic nerves of anesthetized and immobilized animals were imaged using a spinning disc confocal microscope (Fig. 1B, and Mov. 1) at 1 Hz for 1 min, to measure axonal mitochondria movement through kymographs (Fig. 1C and 1D). To determine the stability of Mitotracker labeling over time, optic nerves were imaged 3.5 or 18 hrs after intravitreal Mitotracker injection. Since the distribution (Fig. S1A) and velocity (Fig. 1SB) of stationary, anterogradely-moving, and retrogradely-moving mitochondria were similar at both timepoints, all subsequent imaging experiments were carried out 15-18 hours after Mitotracker intravitreal injection. The kymographs of axonal RGC mitochondria were used to determine the fraction of mitochondria that were stationary, defined here as moving less than 0.1 μm/s, as opposed to moving anterogradely and retrogradely (Fig. 1C and 1D). In an analysis of over 500 axonal mitochondria derived from 6 animals we found that stationary mitochondria account for nearly half of total axonal mitochondria whereas both anterogradely and retrogradely transported mitochondria each contribute for a quarter of the total (Fig. 1E), consistent with other studies of axonal mitochondria (Zheng et al., 2019; Fellows et al., 2020; Suh et al., 2021). Amongst the motile mitochondria, the average speed was 0.63 μm/s and 0.78 μm/s for anterogradely and retrogradely moving mitochondria, respectively (Fig. 1F), which is within the range of the values previously measured by others (Morris and Hollenbeck, 1993; Ligon and Steward, 2000; Vos et al., 2003; Miller and Sheetz, 2004; Jiménez-Mateos et al., 2006; Pilling et al., 2006; Mironov, 2007; Misgeld et al., 2007; Kang et al., 2008; Wang and Schwarz, 2009; MacAskill and Kittler, 2010; Chang et al., 2011; Chen et al., 2016; Niescier et al., 2016). Thus, the *X. laevis* tadpole optic nerve was deemed a suitable model to study RGC axonal mitochondria.

**Figure 1.**
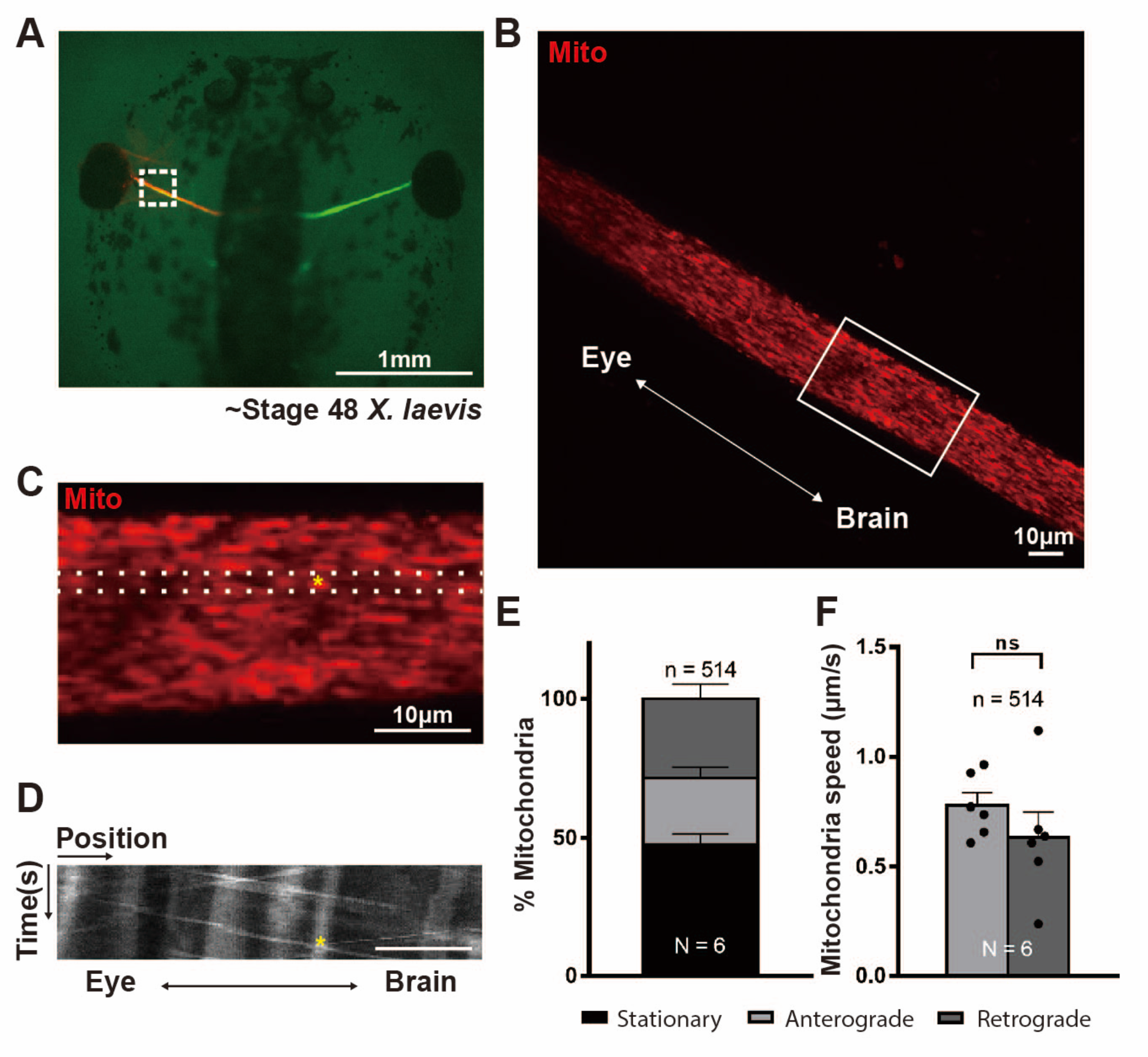
Live-imaging mitochondria movement within *X. laevis* tadpole optic nerve shows that in RGC axons approximately half of mitochondria are stopped and there is balanced anterograde and retrograde movement. (**A**) Intravitreal injection of Mitotracker Deep Red can be used to label RGC mitochondria within axons; here in *Tg(Isl2b:GFP)* transgenic tadpoles where the RGC axons are also labeled by a cytoplasmic GFP expressed by the *Isl2b* RGC-specific promoter. The white dotted box illustrates the approximate location of spinning-disc confocal live-imaging (1 min, 1 Hz). Representative single frame of a Mitotracker Deep Red labeled optic nerve (full region imaged in **B**; boxed area rotated and enlarged in **C**). (**D**) Representative kymograph displaying the position (*x* axis) of mitochondria over time (*y* axis, 60s); derived from the region between the dotted lines in C for all 60 frames. The yellow star indicates the position of the same mitochondrion in both kymograph (**D**) and representative image (**C**). Scale bar, 10 µm. (**E**-**F**) Quantification of mitochondrial movements, showing that: (**E**) About half of RGC axonal mitochondria are stationary and there are comparable numbers of mitochondria moving anterogradely and retrogradely, and (**F**) the average speed of anterograde and retrograde mitochondria movement are similar. Mean ± SEM; n = 514 mitochondria from 6 animals. Statistical analysis in **F** was performed by unpaired, two-tailed Student’s t-test. Not significant (ns).

Next, to test whether the Mitotracker intravitreal injection itself might affect mitochondria movement within the optic nerve, we generated transgenic animals to image RGC axonal mitochondria without the use of Mitotracker. By crossing of *Tg(Isl2b:GFP)* animals with *Tg(Isl2b:Tom20-mCherry)* animals, we obtained doubly transgenic animals which express both a mitochondria-targeted mCherry fluorescent reporter as well as a cytoplasmic GFP transgene, both specifically in RGCs. Mitotracker Deep Red was injected into *Tg(Isl2b:GFP)* expressing tadpoles at NF stage 48, into both animals with or without expression of the Isl2b:Tom20-mCherry transgene, and then the optic nerves were live-imaged as described above within one day of the intravitreal Mitotracker injection. Most of the mCherry and Mitotracker objects colocalized with each other both in the merged images (Fig. S1C) and kymographs (Fig. S1D), indicating that the mitochondria-targeted transgene and Mitotracker similarly label the RGC axonal mitochondria. The Mitotracker-based quantifications showed no significant difference in mitochondria movement between the two different Mitotracker injected groups (with and without presence of the mitochondria-targeted transgene), whereas the Tom20-based quantifications showed that the Mitotracker intravitreal injection does have a small but significant effect on slowing retrograde transport and increasing the fraction of stopped mitochondria at the expense of orthograde moving mitochondria (Fig. S1F and S1E, respectively). Because of this effect, in all subsequent experiments, animals with or without intravitreal injection were compared only to other animals similarly treated.

### Doxycycline-induced expression in RGCs of disease-associated OPTN mutants leads to changes in movement and co-localization indicative of increased axonal mitophagy within the optic nerve

Mutations in OPTN can lead to impaired or aberrant mitophagy and the accumulation of defective mitochondria (Wong and Holzbaur, 2014a; Lazarou et al., 2015; Shim et al., 2016; Evans and Holzbaur, 2020). To test whether OPTN mutations might affect the behavior of RGC mitochondria and components of the mitophagic machinery, OPTN and LC3b, specifically within axons, we generated transgenic *X. laevis* in which fluorescently tagged LC3b and variants of OPTN that could be expressed selectively in RGCs by induction with doxycycline (Fig. 2A and 2B). In these large transgene constructs, an Isl2b RGC-specific promoter drives a version of the reverse tetracycline trans-activator, rtTA2, that is highly efficient in *X. laevis* (Das and Brown, 2004; Mills et al., 2015) in combination with a bi-directional tetracycline operator that expresses EGFP-LC3b in one direction and in the other mCherry-tagged versions of the following human OPTN variants: Wt, glaucoma-associated mutations E50K, M98K and H486R, ALS-associated mutation E478G, and two synthetic mutations, F178A and D474N, previously shown to be defective in the OPTN interaction with LC3b and ubiquitinated mitochondria, respectively (Wild et al., 2011; Korac et al., 2013; Wong and Holzbaur, 2014a; Sirohi et al., 2015; Shim et al., 2016; Li et al., 2018; Liu et al., 2018; Chernyshova et al., 2019; Padman et al., 2019). The optic nerves of 4-6 transgenic tadpoles per transgene were imaged using the spinning-disc confocal first as a 1- minute single plane t-series acquired at 1 Hz, and then as a z-scan imaging the full thickness of the optic nerve at 1 µm steps. In the case of animals expressing E50K and Wt OPTNs, analyses were done with and without previous labeling of RGC mitochondria through intravitreal injection of Mitotracker (Fig. 2C and 2D); in the case of all other OPTN variants analyses of OPTN and LC3b were carried only without Mitotracker labeling in order to avoid the possible confounder introduced by the intravitreal Mitotracker injection. Kymograph analyses of the t-series were used to quantify the percentage of each movement class (stationary, anterograde and retrograde) and determine the average speed of anterograde and retrograde movement. For the live-imaging and kymograph analyses of OPTN and LC3b, in the absence of Mitotracker intravitreal injection, analyses were carried out in both primary transgenic F0 animals (Fig. 2E, 2E’, S2C and S2D) as well as in their F1 progeny (Fig. S2A, S2B, S2E and S2F), to minimize the possibilities of transgene copy number or integration position effects confounding the interpretation. Such analyses in F0s are possible in *X. laevis* because the restriction enzyme mediated integration (REMI) transgenesis method used to generate the animals (Kroll and Amaya, 1996) involves transgene integration prior to the first cell division, and thus provides fully transgenic rather than mosaic animals. The results from F0 transgenic founders and F1 progeny were very similar and showed that relative to Wt OPTN, all OPTN mutants with the exception of the LC3b-binding-defective F178A mutant trended towards or had significant increases in the fraction of both stopped OPTN and LC3b, with the effects being largest for the glaucoma-associated mutations E50K or M98K. Analyses of antergrade and retrograde velocity for OPTN and LC3b in both F0s (Fig. S2C and S2D) and F1 animals (Fig. S2E and S2F) found no consistent significant changes in velocity, suggesting the increases in stopped OPTN and LC3b were specific effects and not secondary to making the RGCs or their axons unhealthy. To determine whether these increases in stopped OPTN and LC3b might represent an increase in axonal mitophagy, the studies of OPTN and LC3b movement were repeated, but now 15 to 18 hrs after intravitreal injection of Mitotracker (Fig. 2F); these studies were carried just in F1 animals expressing Wt or E50K OPTN, as well as in animals carrying no transgenes at all. Compared to the non-transgenic control animals (only Mitotracker-injected) those with expression of Wt OPTN had no alteration in the fraction of stopped mitochondria, similar to what had been previously observed after expression of the Tom20-mCherry transgene (Fig. S1E). In contrast, expression of E50K OPTN resulted in a significant increase in the fraction of stopped mitochondria, above the already high baseline of stopped mitochondria (Fig. 2F). Of note, as was the case for OPTN and LC3b in the absence of intravitreal Mitotracker injection, expression of neither Wt or E50K OPTN affected the relative balance of retrograde versus orthograde movement of mitochondria, OPTN or LC3b, or their velocities, but rather only increased the fraction of the stopped populations. Consistent with the previous results showing that intravitreal injection of Mitotracker had a small but significant effect on mitochondria movement (Fig. S1E and S1F), here too we found that the intravitreal injection of Mitotracker affected the behavior of axonal OPTN, including an increase in the fraction of stopped OPTN; however, so did animals receiving an injection of just the biological salts solvent, showing that it is the eye injection procedure rather than the Mitotracker itself that mainly perturbs the system (Fig. S2H).

**Figure 2.**
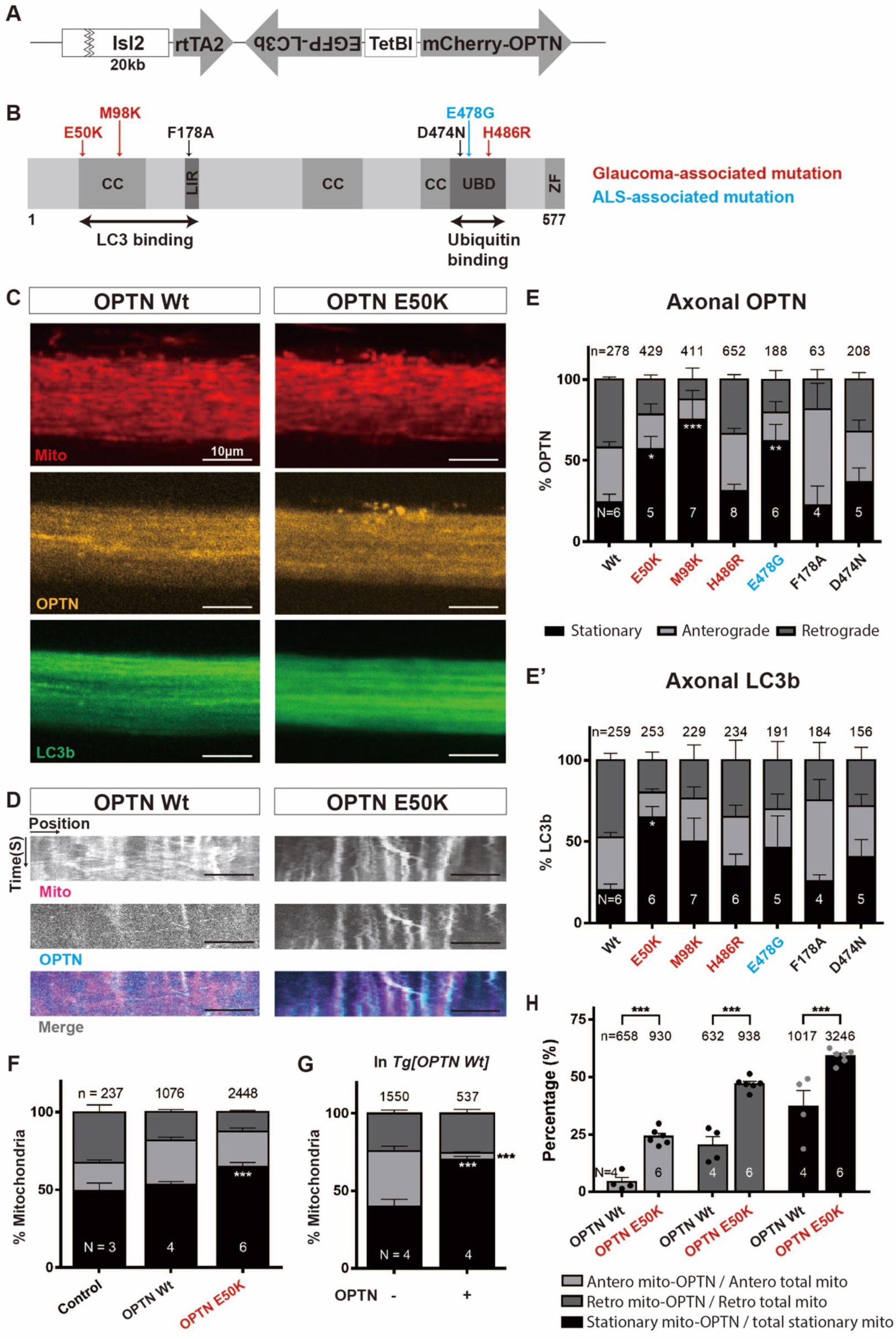
OPTN mutants conditionally expressed in RGCs increase the fraction of stopped axonal mitochondria, OPTN and LC3b and the fraction of mitochondria co-localizing with OPTN. (**A**) Illustration of the transgenic construct: Isl2b promoter driving rtTA2 linked to EGFP-LC3b and mCherry-OPTN driven in opposite strands by a bidirectional tetracycline operator, pTRETightBI. The fusion-constructs are thus expressed only in RGCs and only after doxycycline induction. (**B**) Schematic of OPTN functional domains showing the position of the point mutations examined. Three glaucoma-associated mutations, E50K, M98K and H486R; An ALS-associated mutation, E478G; Two synthetic OPTN mutations, F178A and D474N, disrupting LC3b and ubiquitin binding, respectively. CC: coiled coil domain, LIR: LC3 interacting region, UBD: ubiquitin-binding domain, ZF: zinc finger. Numbers indicate the amino acid position. (**C**-**D**) Representative confocal images (**C**) and corresponding kymographs (**D**) of axonal mitochondria, OPTN and LC3b in Mitotracker-injected *Tg(Isl2b:mCherry-OPTN(Wt or E50K)_ EGFP-LC3b)* animals three days after induction of transgene expression. Scale bar, 10 µm. (**E**-**E’**) Quantification of OPTN (**E**) and LC3b (**E’**) movements in Wt OPTN and various OPTN mutants (all independent F0 animals). Bar graphs show the percentage of each movement (stationary, anterograde and retrograde) of OPTN and LC3b in Wt OPTN and OPTN mutants. Mean ± SEM; n = 63-652 OPTN and n = 156-259 LC3b objects measured in 4-8 animals. (**F**) Quantification of mitochondrial movements after Mitotracker injection in control (non-transgenic), and transgenic animals expressing in RGCs either Wt or E50K OPTN (analyzed in F1 animals). Expression of E50K OPTN but not Wt OPTN results in a significant increase in stalled mitochondria compared to the control. Mean ± SEM; n = 237-2448 mitochondria from 3- 6 animals. (**G-H**) Quantification of mito-OPTN colocalization in Mitotracker-injected Wt OPTN (**G, H**) and E50K OPTN (**H**) animals. (**G**) Percentage of each movement (stationary, anterograde and retrograde) of mitochondria co-localizing (mito-OPTN) or not co-localizing (mito-ONLY) with OPTN in the animals expressing Wt OPTN. The majority of mito-OPTN objects are stationary, whereas mitochondria not co-localizing with OPTN have balanced stationary, anterograde and retrograde movement. Mean ± SEM; n = 1550 mito-ONLY, n = 537 mito- OPTN colocalizations from 4 Wt OPTN animals. (**H**) Expression of E50K OPTN results in increased fraction of mito-OPTN (mitochondria-OPTN colocalization) relative to Wt OPTN, both in the moving and stationary pools. Mean ± SEM; n = 658-930 total orthograde mitochondria, n = 632-938 total retrograde mitochondria, n = 1017-3246 total stationary mitochondria from 4-6 animals. Statistical analysis in **E**-**H** were performed by two-way ANOVA following Tukey’s post-hoc test for multiple comparisons. * = p<0.05, ** = p<0.01, *** = p<0.001.

Next, to determine whether the increases in stopped mitochondria and stopped OPTN might indicate an increase in axonal mitophagy, the degree of mitochondria and OPTN co- localization was examined. In the nerves labeled by Mitotracker, visual inspection of the raw images (Fig. 2C) and the derived kymographs (Fig. 2D) showed that OPTN and the Mitotracker-labeled mitochondria often co-localized, particularly in the stopped populations, and more so in the animals expressing E50K OPTN, further suggesting that at least a fraction of the stopped LC3b, OPTN and mitochondria might represent mitophagy occurring in the axons. To further define the relationship between stopped objects and mitophagy, we re-analyzed the previous kymographs of mitochondria and OPTN and separately quantified mitochondria that co-localized with OPTN (mito-OPTN) versus mitochondria not associated with OPTN (mito-ONLY) (Fig. 2G and S2I). Strikingly, while mitochondria not colocalizing with OPTN had relatively balanced populations of stopped, anterograde and retrograde movement, the majority of mitochondria colocalizing with OPTN were stopped, with the remainder largely being engaged in retrograde transport; the same was true in animals expressing Wt OPTN (Fig. 2G) and E50K OPTN (Fig. S2I). The glaucoma-associated E50K OPTN also significantly increased the fraction of mitochondria co-localizing with OPTN, and that was evident in both the stationary and the two moving populations (Fig. 2H). In summary, expression of OPTN versions carrying glaucoma and ALS-associated mutations, as well as a synthetic mutation in OPTN that perturbs the recognition of ubiquitinated mitochondria, but not a synthetic mutation that perturbs the association of OPTN with LC3b, all lead to increases in stopped axonal OPTN and LC3b in RGCs. At least in the case of E50K OPTN, which increases OPTN-mitochondria colocalization, some of these stopped OPTN likely represent mitochondria undergoing mitophagy or something akin to mitophagy locally within the optic nerve.

### Expression of the glaucoma-associated E50K OPTN leads to increased accumulation of mitochondria and OPTN outside of axons

In many of the animals, particularly those expressing E50K OPTN, some of the mitochondria and OPTN signal appeared to be not co-localized with the LC3b signal, but rather to be on the surface of the optic nerve (e.g., middle top of the OPTN E50K nerve shown in Fig. 2C). To quantify such extra-axonal mitochondria and OPTN, the Z-scans of the full thickness of the optic nerves, acquired at 1 µm steps, were analyzed using the 3D imaging software Imaris. Because LC3b localizes not only to autophagosomal membranes, but also is in the cytoplasm/axoplasm (Xie and Klionsky, 2007; Mizushima and Komatsu, 2011; Fu et al., 2014; Wong and Holzbaur, 2014b; Tammineni et al., 2017; Stavoe et al., 2019; Evans and Holzbaur, 2020; Boecker et al., 2021; Kuijpers et al., 2021), we used the EGFP-LC3b signal to create a mask to demark the RGC axons, so as to then be able to separately measure the OPTN (mCherry) or mitochondria (far-red) signals that co-localized or not with the axons. First, such analyses were carried out for the same animals above analyzed for OPTN and LC3b movement metrics, but that had not been subjected to the intravitreal injections of Mitotracker, animals in which the expression of mCherry-hOPTNs (Wt, E50K, M98K, F178A, D474N, E478G and H486R) and EGFP-LC3b was induced in RGCs through bath application of doxycycline. Measurements carried out in the 3D reconstructions showed significant increases in the fraction of OPTN outside of the RGC axons in the optic nerves of glaucoma-associated mutations, especially E50K (12.6%), when compared to Wt OPTN (0.5%), ALS-associated mutation E478G (0.2%), and the synthetic mutations F178A (3.3%) and D474N (0.1%) (Fig. 3B). Then, to determine whether such extra-axonal OPTN might be related to the degradation of mitochondria, the same procedure was applied to the Z-scan images of animals where Wt OPTN and E50K OPTN were similarly induced by doxycycline, but that also a day prior had received an intravitreal injection of Mitotracker (Fig. 3C). Imaris-based 3D reconstruction and volume quantification revealed that expression of OPTN carrying the glaucoma-associated E50K mutation results in a significant increase in both the fraction of all stopped RGC axonal mitochondria outside of the LC3b-labeled RGC axons (13.4%) but also the fraction of all stopped RGC axonal mitochondria co-localizing with OPTN also outside of the LC3b-labeled RGC axons (17.9%), as compared to the much lower amounts found after expression of Wt OPTN (0.5% for both mitochondria and mito-OPTN) (Fig. 3D and 3E, and Mov. 2). This fractional value likely relates to only the stopped mitochondria, as the z-scan imaging used frame-averaging, and moving objects were blurred and likely underrepresented. In summary, expression of E50K OPTN, but not OPTN that lacks the glaucoma-associated mutation, not only increases the amount of stopped mitochondria, OPTN and LC3b, and the degree of colocalization between mitochondria and OPTN within axons, all of which can be interpreted as indications of axonal mitophagy, but also results in large amounts of mitochondria and the mitophagy receptor OPTN being found outside of the axons.

**Figure 3.**
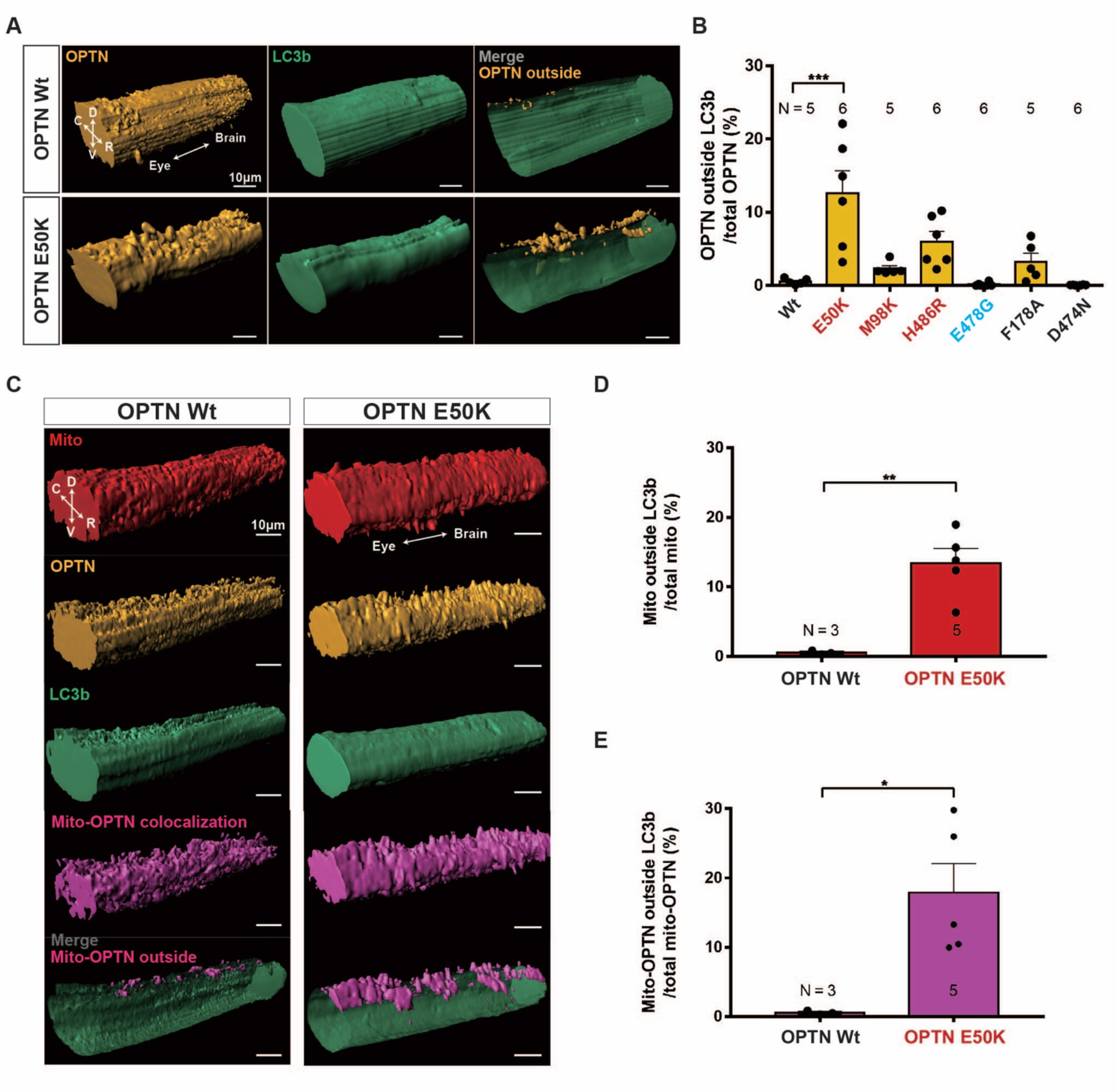
3D reconstruction of axonal mitochondria, OPTN and LC3b in optic nerves shows that glaucoma-associated OPTN mutations increase the amounts of mitochondria, OPTN and mito- OPTN outside of the LC3b-labeled axons. (**A**-**B**) (**A**) Reconstruction of optic nerves displaying OPTN, LC3b and OPTN outside LC3b-labeled RGC axons (merged images on the right) in the optic nerves of animals expressing Wt and E50K OPTN. D: dorsal, V: ventral, C: caudal, R: rostral. Scale bar, 10 µm. (**B**) Percentage of OPTN outside LC3b-labeled RGC axons is significantly higher or trends so in the glaucoma-associated OPTN mutations. Mean ± SEM; n = 5-6 animals. (**C**-**E**) 3D reconstruction of axonal mitochondria, OPTN and LC3b within the segments of intraocular optic nerves in Mitotracker-injected *Tg(Isl2b:mCherry-OPTN(Wt or E50K)_ EGFP-LC3b) X. laevis*. (**C**) Reconstruction displaying mitochondria, OPTN, LC3b, mito-OPTN colocalization and mito-OPTN outside LC3b-labeled RGC axons (merged images on the bottom) in the optic nerves of animals expressing Wt and E50K OPTN. D: dorsal, V: ventral, C: caudal, R: rostral. Scale bar, 10 µm. (**D**-**E**) Percentage of mitochondria (**D**) and mito-OPTN (**E**) outside of LC3b-labeled RGC axons after expression of Wt and E50K OPTN. Expression of E50K OPTN results in a significant increase in mitochondria and mito-OPTN colocalizations outside of LC3b- labeled RGC axons. Mean ± SEM; n = 3-5 animals. Statistical analysis in **B** was performed by one-way ANOVA following Tukey’s post-hoc test for multiple comparisons, and **D**-**E** were performed by unpaired, two-tailed Student’s t-test. * = p<0.05, *** = p<0.001.

### Sparse labeling of axons shows even larger amounts of axonal mitochondria and OPTN outside of axons and that only in the most extreme cases are they observed on the ON surface

In the experiments just described, the majority of the LC3b-negative-OPTN-positive mitochondria signal was found on what appears to be the surface of the optic nerve, which was most obvious after expression of E50K OPTN (see Fig. 3A and C, and Mov. 2). To determine whether additional extra-axonal OPTN and mitochondria might be found within the optic nerve parenchyma but obscured by the EGFP-LC3b mask due to the high axon density and low imaging resolution, we turned to a sparse labeling approach. Small groups of retinal cells, estimated to be 10-50 in number, were surgically transplanted from E50K OPTN transgenic progeny into progeny of a line expressing *Tg(Fabp7:mTagBFP2-Ras)*, in which astrocyte membranes are labeled by a membrane targeted fluorophore expressed by a promoter from the fatty acid binding protein 7 gene (*fabp7*, also known as basic lipid binding protein or *blbp*) (Owada et al., 1996; Sharifi et al., 2011) which drives expression in frog optic nerve astrocytes (Mills et al., 2015). Transplants were done around NF stage 24-32, when all cells in the retina are progenitors and thus prior to RGC differentiation. A week later in such animals and 3-days after doxycycline induction, small groups of axons were readily visible in these optic nerves based on the EGFP-LC3b signal. 3D reconstruction and volume measurements of the astrocyte membrane signal derived from the host and the OPTN and LC3b signals derived from E50K OPTN expressing donor cells (Fig. 4A and 4B, and Mov. 3) showed that the fraction of OPTN signal on the surface of the optic nerve was 17.1% of the total signal, in relatively good agreement with the data previously obtained in the whole nerves (see Fig. 3B). However, this represented only 23.9% of the total OPTN outside of the LC3b-labeled axons, the rest being within the optic nerve parenchyma. Furthermore, the nerves with the largest amounts of OPTN on the surface of the optic nerve were those with the greatest amount of extra-axonal OPTN (Fig. 4B). These sparse labeling data thus suggested that our previous measures of the amount of OPTN and mitochondria outside of axons quantified in Fig. 3 based on the whole-nerve imaging after expression of all OPTN variants likely grossly underreported the actual amounts of OPTN and mitochondria outside of the axons. Thus, for animals expressing either Wt or E50K OPTN, we repeated the same t-series and z-scan imaging and following kymograph analyses and Imaris 3D reconstructions in the context of the sparse labeled axons, but also after dye-labeling the mitochondria. One hour after injection of Mitotracker into the eye anlage, retina progenitor cells from Wt or E50K OPTN transgenic progeny were transplanted into the eyes of non-transgenic hosts (Fig. 4C, S3A and S3B). Consistent with our previous data, more stalled mitochondria, OPTN and LC3b puncta were observed in the axons expressing E50K OPTN compared to those expressing Wt OPTN, with relatively balanced populations of orthograde and retrograde movement in both Wt and E50K OPTN (analyzed per axon; Fig. S3C and analyzed per animal; Fig. S3C’). Orthograde and retrograde velocities for mitochondria, OPTN and LC3b in the sparsely labeled Wt and E50K OPTN axons were similar to those measured in the whole nerves, and once again showed no significant differences in velocities after expression of E50K OPTN (Fig. S3D). Notably, however, the amounts of mitochondria and OPTN outside of axons were indeed higher than those previously measured. In the case of E50K OPTN, 35.8% of mitochondria and 21.8% of OPTN were determined to be outside of the axons in the sparse labeling experiments (as compared to 13.4% and 17.9%, respectively, measured in the whole nerves). The difference between whole nerve and sparsely labeled axons was far larger in the case of expression of Wt OPTN, where previously approximately 0.5% had been found for both (see Fig. 3D and 3E), but now 15.2% of mitochondria and 5.7% of OPTN were shown to be outside of the RGC axons (Fig. 4D and 4E); this order of magnitude difference in results is presumably due to in the case of the presence of Wt OPTN nearly all the extra-axonal mitochondria and OPTN being found within the nerve parenchyma, not having reached high enough levels to be measurable on the surface of the optic nerve. Collectively, these data show that even in the case of expression of Wt OPTN, which we showed above does not in any way alter the movement behavior of mitochondria within axons, there already is a substantial amount of both mitochondria and OPTN outside of the axons from which they derive, and that expression of E50K OPTN, in addition to resulting in the stalling of large number of mitochondria, OPTN and LC3b within axons and colocalization of mitochondria and OPTN, also significantly increases the amount of both OPTN and mitochondria that are found outside of the axons.

**Figure 4.**
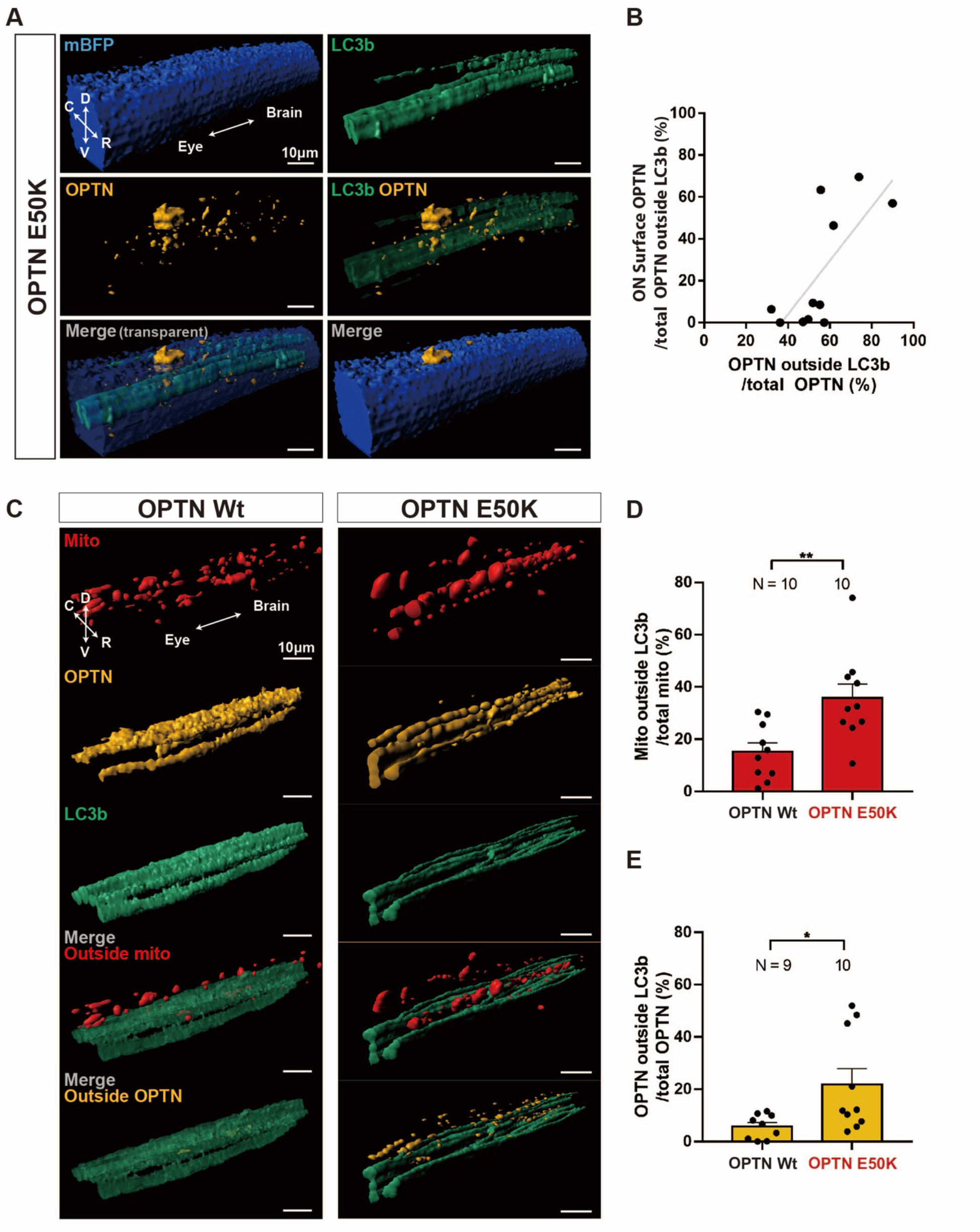
The RGC axonal mitochondria and OPTN found outside of the LC3b-labeled axons are both within the optic nerve parenchyma and on the surface of the optic nerve, and are increased by expression of E50K OPTN. (**A**-**B**) E50K OPTN from sparsely labeled axons found outside of LC3b-labeled axons are found both within optic nerve parenchyma and optic nerve surface. (**A**) Reconstruction showing astrocyte membranes of the host, from *Tg(Fabp7:mTagBFP2-Ras)* here abbreviated as mBFP, and E50K OPTN and LC3b (merged images on the bottom) from the axons of the donor cells. D: dorsal, V: ventral, C: caudal, R: rostral. Scale bar, 10 µm. (**B**) Percentage of OPTN on the optic nerve surface (y-axis) vs. percentage of total OPTN outside of the LC3b-labeled axons (x-axis). n = 11 animals. Trend line with 0.522 R-squared value. (**C**-**E**) Sparse labeling of axons reveals that extensive amounts of axonal mitochondria and OPTN are outside of the axons. (**C**) 3D-reconstructions of sparsely labeled axons showing mitochondria and either Wt or E50K OPTN reveal much of both are outside the LC3b-labeled RGC axons (merged images on the bottom). D: dorsal, V: ventral, C: caudal, R: rostral. Scale bar, 10 µm. (**D**-**E**) Percentage of mitochondria (**D**) and OPTN (**E**) outside the LC3b-labeled RGC axons in animals expressing Wt and E50K OPTN, respectively. Mean ± SEM; n = 9-10 animals. Statistical analysis in **D** and **E** were performed by unpaired, two-tailed Student’s t-test. * = p<0.05, ** = p<0.01.

### Shed axonal mitochondria are degraded by astrocytes

To determine where outside axons the axonal mitochondria were degraded, we carried out correlated light-EM analyses on sparsely labeled axons. First, cells from animals expressing a membrane-associated mCherry transgene in RGCs were transplanted into animals expressing two different transgenes in astrocytes, the same membrane-associated BFP transgene used above to label all astrocyte membranes, but also an Aquaporin-4-GFP (Aqp4-GFP) fusion construct that labels principally the glial limitans at the surface of the optic nerve. One day after labeling the RGC axonal mitochondria through an intravitreal injection of Mitotracker Deep Red, the nerves of these animals were live-imaged in their entirety at lower x,y and z-resolution for just the Aqp-GFP transgene (Fig. 5A), and then one region imaged at higher resolution in all four channels (Fig. 5B). The heads of fixed animals, heavy metal processed and embedded in epoxy resin, were analyzed by micro-CT in order to identify the same region that had been live imaged (Fig. 5C) and then the full thickness of the optic nerve at that location was analyzed by serial block-face scanning electron microscopy (SBEM). After registering the live imaging and SBEM datasets based on the position of nuclei, some of the stopped mitochondria that had been live-imaged could be unambiguously identified at the level of EM (Fig. 5D-F). The axon-derived material on the surface of the optic nerve, in this case dye-labeled mitochondria and transgene-labeled axolemal membranes, were determined to be within the soma of the transgene labeled astrocytes (Fig. 5D- E), and morphologically appeared as mitochondria and membranes at various stages of degradation. The mitochondria that live-imaging determined to be stationary within the nerve parenchyma were harder to unambiguously correlate between light and SBEM datasets, but those that could were determined to be within the processes of astrocytes (Fig. 5F). Notably, while axonal debris was most often observed within astrocyte fine processes, often those processes were closely associated with fine processes of distinct astrocytes (Fig. 5F). The soma of the astrocytes were located at the optic nerve periphery but had processes that extended deep into the parenchyma (Fig. 5G), similar to astrocytes that surround axon bundles within larger optic nerves. Thus, it is possible that the axon-derived debris found within astrocyte soma at the surface of the optic nerve might have originated deep within the optic nerve parenchyma. However, it also may have originated near the astrocyte soma, as dystrophic axons with mitochondria undergoing early steps of mitophagy could also be observed within some axons in close contact with the astrocyte soma (Fig. 5H-I).

**Figure 5.**
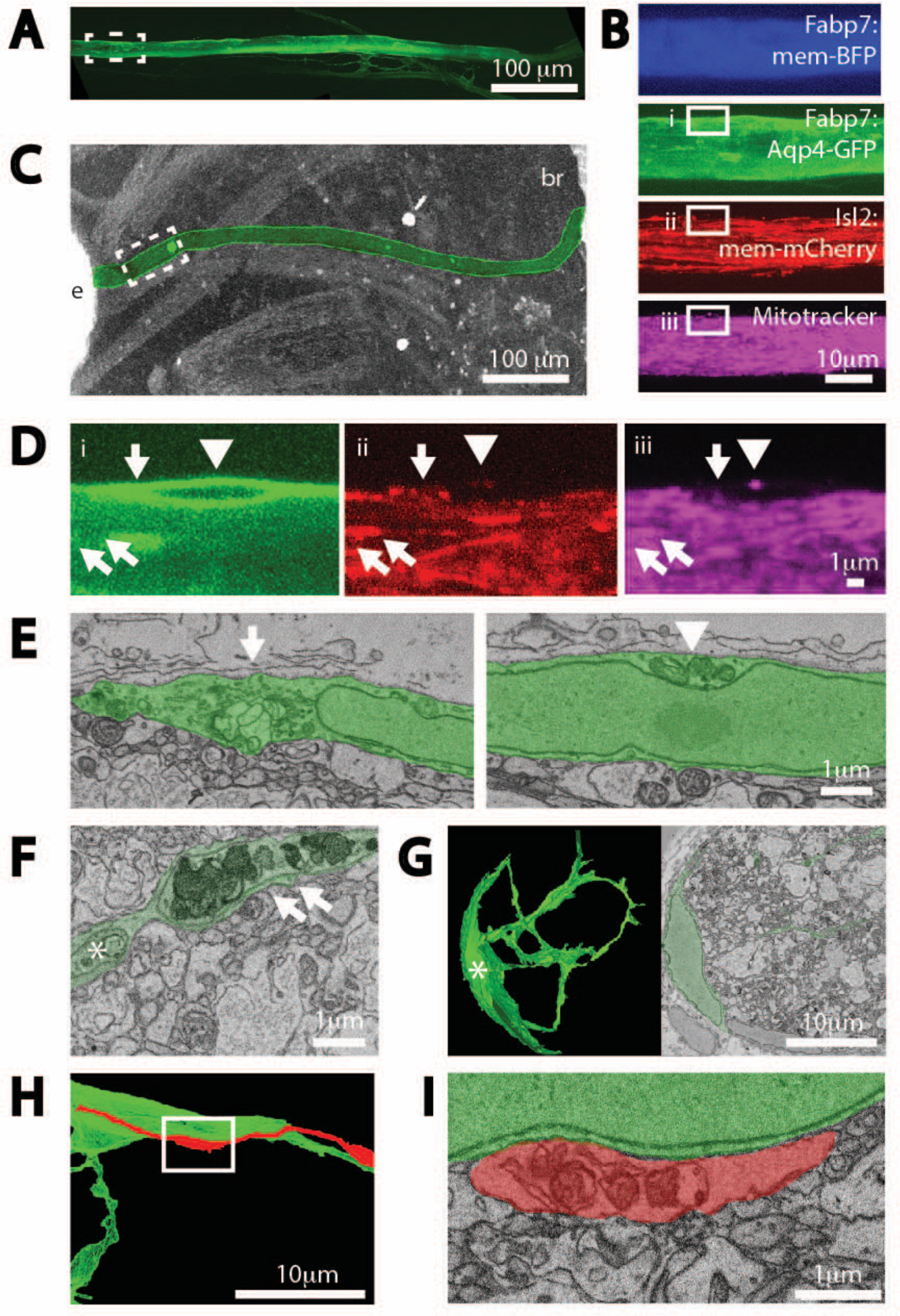
Extruded axonal mitochondria are degraded by astrocytes. (**A**) Z-projection of full length of the optic nerve visible through live imaging, labeled by an Aqp4-GFP transgene expressed in astrocytes. **(B)** Z-projections of four channels after higher resolution imaging of a region of the same optic nerve, position shown in stippled boxed region in A; four channels are astrocyte expressed mem-BFP and Aqp4-GFP transgenes, an RGC expressed mem-mCherry transgene, together with intravitreally injected Mitotracker. Solid boxes represent fields shown at higher resolution in D-E. (**C**) Micro-CT of the head of same animal after embedding in epoxy-resin, used to pin-point the area live-imaged for subsequent SBEM analyses. Optic nerve is colorized green. Letter insets: e is eye and br is brain. **(D)** Superficial astrocyte soma labeled by astrocyte expressed Aqp4-GFP transgene (i) containing axonal derived membranes (ii) and mitochondria (iii). Arrow, arrowhead and double arrows represent three discrete axonal mitochondria signal unambiguously identifiable at both light level live-imaging, but also at the level of SBEM, shown in E-F. **(E)** Two SBEM sections of the same astrocyte soma centered on discrete pockets phagocytosed axonal membranes and mitochondrial material. **(F)** Extra-axonal mitochondria and membranes within fine astrocyte processes. Asterisk represents a process originating from same astrocyte shown in G. **(G)** Morphology of one the astrocytes whose processes are near the extra-axonal mitochondria shown in F; on right, reconstruction of about half the astrocyte and on left single plane. **(H)** Reconstruction of an axon near the soma of the same astrocyte shown in D-E. **(I)** Dystrophic mitochondria within axon appearing to be enwrapped in other axonal membranes within the axon, adjacent to superficial astrocyte soma.

### OPTN and mitochondria leave RGC axons in the form of exophers

To investigate how RGC axonal OPTN might leave axons within the optic nerve parenchyma, we performed repetitive z-scan live-imaging of sparsely labeled axons for 5 minutes at 30-second intervals. Since the shedding intermediates were presumed to be rare and short-lived, these studies were conducted mainly for the M98K OPTN glaucoma-associated mutation, as it appeared more severe than the E50K OPTN mutation at least in the fraction of stopped OPTN in both F0 and F1 analyses (Fig. 2E and Suppl Fig. 2A). After the repetitive z-scan live-imaging, areas that had focally increased LC3b and OPTN signal at the sites of focal changes in axonal diameter were digitally extracted and their position at different times registered, so as to measure shape and fluorescence intensity changes over time. The four types of dystrophies observed were symmetric axonal swellings, asymmetric axonal dystrophies with static or changing levels of OPTN or LC3b, asymmetric axonal dystrophies that are pinched off from the axon during the imaging window, and structures with no connection to axons. Examples of only the first three are here shown, as the fourth provided no mechanistic insight as to how they might become separated from the axons. An axonal “swelling” is shown where the mitophagy machinery OPTN and LC3b are focally stopped and enriched together (Fig. 6A-i). An asymmetric axon dystrophy (i.e., axonal “protrusion”) is shown in what appears to be a “loading” stage (Fig. 6A-ii and Mov. 4). Here, the OPTN signal increases within the exopher-like dystrophy simultaneously with a decrease in the OPTN signal immediately below the dystrophy within the associated axon, suggestive of an active loading process (Fig. 6B). At least some of these asymmetric dystrophies appear to become separated from their axon of origin. The last example is a “pinching-off” event (i.e., “evulsion”) where an exopher-like structure is physically detached from the axon of origin and appears to amalgamate with other areas of extra-axonal OPTN; these are presumed to be within acidified organelles of the phagocyte as they had no measurable GFP signal (Fig. 6A-iii and Mov. 5). Minimum distance measures between the outer boundaries of exopher-like structure and axon of origin (Fig. 6C) and measures of the distance from the exopher centroid to the nearest position along the axon (not shown) demonstrate that the detachment process is acute, occurring within a 30-second interval. We also observed similar axonal dystrophies and exopher-like structures in E50K OPTN under similar imaging settings, but with 2-min intervals and additional Mitotracker labeling (Mov. 6), demonstrating that these structures not only contain OPTN but also mitochondria or mitochondria remnants. The shape, size and dynamic changes in these focal axonal exophers lead us to conclude that at least some of the OPTN and mitochondria leave the axons by the process we previously described in the mouse optic nerve head, transmitophagy (Davis et al., 2014).

**Figure 6.**
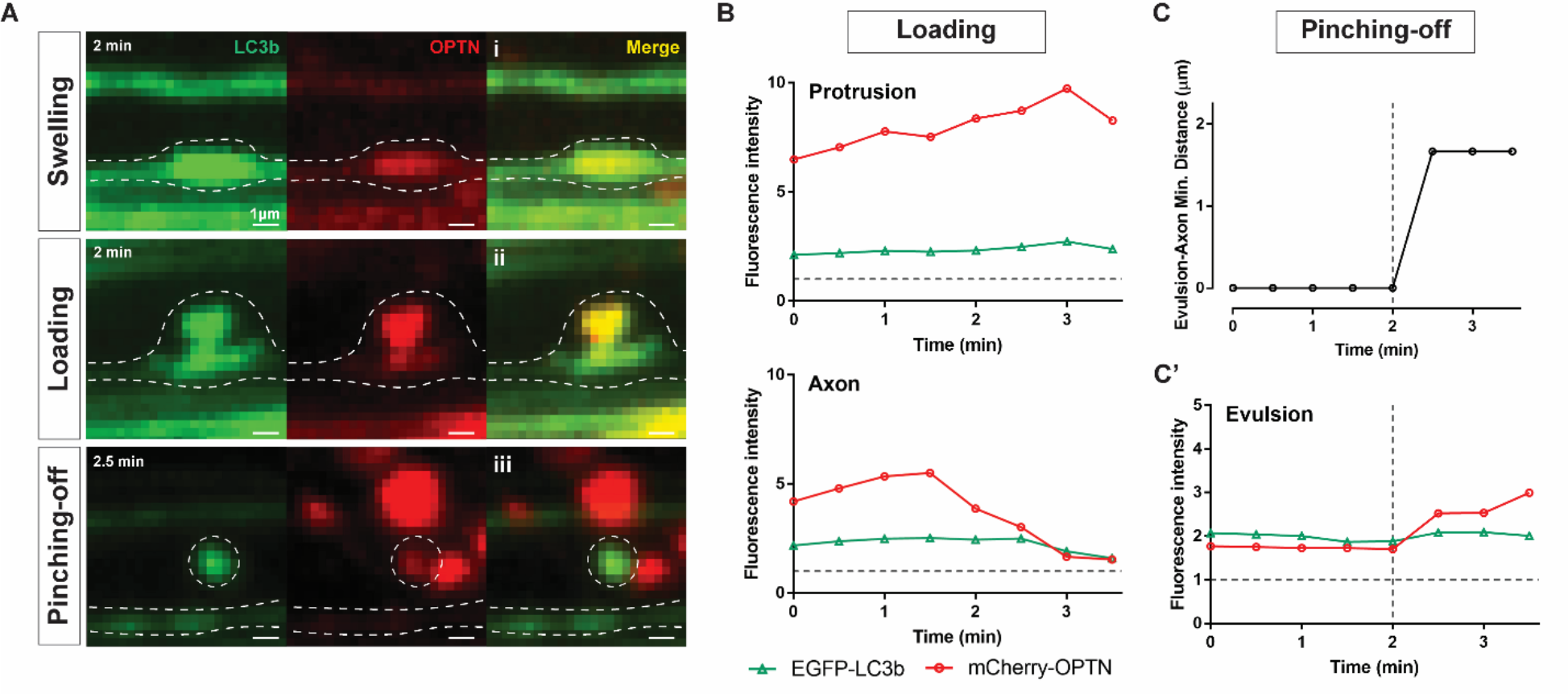
At least some OPTN leaves RGC axons in the form of exophers. (**A**) Examples of axonal dystrophies in sparsely labeled *Tg(Isl2b:mCherry-OPTN(M98K)_ EGFP-LC3b)* axons containing partially co-localized stopped OPTN and LC3b; presented as a pseudo-sequence consistent with extra-axonal OPTN being the product of transmitophagy. **(i)** Swelling, where the mitophagy machinery OPTN and LC3b are stopped together and focally enriched in a symmetric axonal dystrophy. **(ii)** Loading, where focally accumulated OPTN is enriched in an exopher-like asymmetric axonal dystrophy. **(iii)** Pinching-off, where an exopher-like structure is acutely separated from the axon of origin and appears to coalesce with other extra-axonal OPTN presumably within acidified organelles of a phagocyte. Scale bar, 1 µm. (**B**) Changes in fluorescence within the exopher-like structure and directly beneath within the axon of origin suggest an active loading process. (**C**) Minimum distance and (**C’**) changes in fluorescence within exopher-like structure show further increase in OPTN soon after separation from the axon. Gray dotted horizonal lines represent the mean fluorescence in the same axon away from the axon dystrophy, set at 1; thus, all fluorescence values shown represent focal concentrations of both LC3b and OPTN signal, suggestive of these being sites of axonal mitophagy.

## Discussion

There is evidence that damaged axonal mitochondria are mainly delivered back to or near the soma for degradation through OPTN-mediated mitophagy, although these studies were conducted mainly in cultured cells (Moore and Holzbaur, 2016; Evans and Holzbaur, 2020). Our studies *in vivo* support the view that this constitutes a principal mechanism for axonal mitochondria degradation in *X. laevis* RGC axons, as even after expression of Wt OPTN there is within axons very little stationary OPTN or LC3b, and very little colocalization of OPTN with LC3b or mitochondria. However, we have found that after expression of various mutated OPTNs, there are significant increases in the amount of OPTN and LC3b stopped within axons, and, in the one case we analyzed in most detail, the glaucoma-associated E50K mutation, also significant increases in the amount of stopped mitochondria much of which co-localized with OPTN. Recent studies have shown that arresting mitochondrial motility, which is mainly driven by degradation of the motor-adaptor protein Miro, is an early stage of mitophagy where dysfunctional mitochondria are stopped and sequestered for subsequent mitophagy through PINK1/Parkin-mediated pathway in both *in vivo* and *in vitro* settings (Wang et al., 2011; Liu et al., 2012; Lovas and Wang, 2013; Ashrafi et al., 2014; Hsieh et al., 2016). Since we demonstrate that the same perturbations stop both OPTN and mitochondria and they colocalize, our data strongly support the view that mitophagy, or at least some steps generally associated with mitophagy, can occur in RGC axons far from the cell body. Of note, while the glaucoma associated mutations E50K and M98K were the most potent and inducing these surrogate markers for local axonal mitophagy, the one ALS mutation tested, E478G, as well as a synthetic mutation meant to interfere with the association of OPTN with ubiquitinated mitochondria, D474N, also promoted the same. Indeed, the only versions of OPTN that did not lead to increased stopped mitophagy machinery were Wt OPTN and a synthetic mutation that interferes with the association between OPTN and LC3b. Thus, under basal conditions, about half of axonal mitochondria are stopped; these likely represent those mitochondria providing energy and other functions locally needed to support diverse cellular processes including but not limited to action potential regeneration. However, after perturbations as slight as the injection of a balanced salt solution into the eye, the population of axonal mitochondria that are stopped within axons can increase. We believe that these are unlikely to be more mitochondria supporting functions normally associated with axonal mitochondria, but rather that this second stopped population of axonal mitochondria are, at least in part, ones that are undergoing mitophagy or at least a process that shares molecular machinery with mitophagy.

The 3D reconstructions of the live-imaged ONs revealed that the glaucoma-associated OPTN mutations were unique in not only increasing the fraction of stopped mitochondria, OPTN and LC3b, and their colocalizations within the RGC axons, all suggestive of axonal mitophagy, but also led to large amounts of axonal mitochondria and OPTN being found outside of the axons, some of which reached the surface of the optic nerve. Indeed, the studies based on sparse labeling of axons, which likely provide the most accurate estimate of the fraction of axonal mitochondria and OPTN outside of axons, suggest that these populations are large. In the case of expression of Wt OPTN, which we believe represents the basal state as we also show that Wt OPTN expression has no effect on the behavior of axonal mitochondria, as much as 15.2 and 5.7 % of the stopped mitochondria and OPTN respectively are outside of the axons, and in the case of expression of the E50K OPTN, those numbers increase to 35.8 and 21.8 %. Of interest, while diverse OPTN mutations increased the fraction of stopped mitochondria within the optic nerve, it was the most common glaucoma-associated OPTN mutation, E50K OPTN, that led to the largest increase in externalized mitochondria. This may be a coincidence or artifact of the experimental system used. However, it also may suggest that the particular mutations in OPTN that promote glaucoma not only perturb mitophagy within axons, but that they may perturb it in such a way that increases the probability that such mitochondria will be degraded non cell-autonomously.

That mitochondria and OPTN are found outside of axons suggests that these axonal mitochondria must be eliminated by the neighboring cells resident in the optic nerve, either the astrocytes which constitute the major local resident population, other resident cell populations such as NG2-cells or microglia, or alternatively by invading myeloid cells, all which are known to have phagocytic capacity. We have previously shown that in the frog optic nerve, it is astrocytes that are the major phagocytes that clear extensive amounts of axonal and myelin debris using well conserved phagocytic machinery, at least during a developmental remodeling event (Mills et al., 2015). Correlated light EM studies of sparsely labeled axons here too demonstrate that the majority of these axonal material, including mitochondria, are degraded by astrocytes, either in their finer processes that interdigitate deep in the nerve parenchyma, or far from the axons within the soma of the astrocytes, which in tadpoles at this stage reside exclusively on the nerve surface. SBEM studies have not yet been carried out on animals where OPTN has been perturbed, so it remains to be determined how the degradation of axonal mitochondria within astrocytes might be perturbed by these manipulations. The finding of large amounts of axonal mitochondria and OPTN on the surface of the optic nerve in astrocytes with prominent localization of an Aqp4-GFP reporter to their membranes is highly reminiscent of the glymphatic-like system that has been described in the mouse optic nerve (Wang et al., 2020); thus, whether mitochondria debris is cleared by a glymphatic pathway maybe should be investigated.

One of the most important questions is how the axonal mitochondria and OPTN reach the outside of axons. Based on the repetitive z-scan live-imaging of sparsely labeled RGC axons, we provide evidence that at least some of the outside OPTN and mitochondria leave the axons by the process of transmitophagy which we first described in the optic nerve head of wildtype mice (Davis et al., 2014). The axonal structures live-imaged in this study are highly similar in size and shape to the mitochondria-filled protrusions still attached to axons and mitochondria-filled evulsions separated from axons that we had described in mice through both a mitochondria targeted tandem EGF-mCherry reporter and SBEM (Davis et al., 2014). The structures are also similar to what others refer to as exophers, both in *Caenorhabditis elegans* (Melentijevic et al., 2017) and in mammalian cardiac tissue (Nicolás-Ávila et al., 2020). In both of these later cases, these cellular extrusions occur constitutively, increase in response to various stressors, and contain protein aggregates and dysfunctional organelles, including mitochondria. Recently, it was been shown that after the pinching off of the exophers, they are processed to smaller “stary night” vesicular structures that likely represent the endolysosomal intermediates on the path to the eventual degradation by lysosomes (Wang et al., 2023); it seems likely that the puncta containing axonal material that we observed within astrocyte processes and soma both at the light at EM level likely are the equivalent of these “stary night”. Thus, we suggest that axonal transmitophagy is an axonal variant of exopher generation, that may have evolved as an alternative mechanism for long projection neurons such as RGCs to deal with more dysfunctional mitochondria or aggregates in distal regions of axons than could be dealt with by the more conventional cell-autonomous processes of somal or axonal mitophagy. What controls the balance between the cell-autonomous and non cell-autonomous axonal degradation mechanisms remains an open question, although the control of mitochondria stalling, perhaps through the regulation of Miro stability by the PINK/Parkin pathway (Wang et al., 2011; Liu et al., 2012; Lovas and Wang, 2013; Ashrafi et al., 2014; Hsieh et al., 2016), is a possibility worthy of further study.

Another question which has not been answered in this study is whether increased transmitophagy may be a cause of RGC degeneration, as the mutations that result in the maximal amounts of distal axon mitophagy and transmitophagy in the *X. laevis* RGC axons are the very same mutations that in humans cause glaucoma. If an increase in transmitophagy does contribute to glaucoma, one of the possible pathogenic mechanisms might be inflammation resultant from the accumulation of immunogenic mitochondria debris outside of the axons, as was shown in the case of mammalian cardiac exophers (Nicolás-Ávila et al., 2020). Regardless of whether or not transmitophagy contributes to the etiology of glaucoma, such a transcellular clearance system might be a mechanism to handle the accumulation of detrimental aggregates and damaged organelles within diverse axons, which would be obvious relevance to a variety of neurodegenerative disorders that affect axonal biology early in their progression.

## Materials and Methods

### Transgenes

pCS2(Isl2b):rtTA2_pTRETightBI_mCherry-OPTNs_EGFP-LC3b: mCherry flanked by additional restriction sites was amplified from pTRETightBI-RY-0 (Addgene plasmid # 31463 ; http://n2t.net/addgene:31463 ; RRID:Addgene_31463) (Mukherji et al., 2011) using CD160 and CD161 and recloned into XmaI and EcoRV sites of the same vector. Full-length human OPTN, from the Mammalian Gene Collection (CloneID 3457195, purchased from OpenBiosystems) was amplified with CD162 and CD163 and placed into MluI and EcoRV sites, creating pTRETightBI:mCherry-OPTN_NLS-EYFP. The β-globin polyadenylation sequence was then amplified from AAV2:MitoEGFPmCherry (Davis et al., 2014) using CD60 and CD61 and inserted in between EcoRV and AatII sites downstream of OPTN. To make the EGFP-LC3b part of the constructs, the Egfp was first amplified from pCS2:mito-EGFP-mCherry (Davis et al., 2014) using CD53 and CD91h and cloned into pTRETight:MitoTimer (Addgene plasmid # 50547 ; http://n2t.net/addgene:50547 ; RRID:Addgene_50547) (Hernandez et al., 2013), with EcoRI and XbaI, creating the construct pTRETight:EGFP. Then, full-length mouse LC3b from OpenBiosystems (CloneID 5319360) was amplified with CD58 and CD59 and cloned into SalI-NheI sites of this construct in order to make pTRETight:EGFP-LC3b. Then, EGFP-LC3b was moved into the previously created pTRETightBI:mCherry-OPTN_NLS- EYFP by cloning it into the EcoRI and XbaI sites, thus replacing the NLS-EYFP. In order to put both EGFP-LC3b and mCherry-OPTN under inducible control, the first step was to create a version of pCS2:rtTA2, previously shown to have low baseline and high inducibility in transgenic *X. laevis* (Das and Brown, 2004; Mills et al., 2015), but with an additional multiple cloning site to facilitate assembly. The rtTA2 itself was amplified from Blbp:rtTA2 (Mills et al., 2015) using CD158 and CD159, and cloned into PmeI and PacI sites of a version of pCS2 vector where the CMV promoter had been previously replaced by 600bp upstream sequences of the Cardiac Actin promoter (as well as a multiple cloning site or MCS containing ZraI, AatII, PmeI and PacI restriction sites), accomplished by amplifying the promoter itself from *X. laevis* genomic DNA using CD156 and CD157. Then, the majority of pTRETightBI:mCherry-OPTN_EGFP-LC3b, excluding the backbone sequences, was placed between the AatII-NotI sites of this pCS2(MCS- CA600):rtTA2, before removing the CA600 promoter itself by digestion with PacI and NotI followed by blunting and self-ligation. pCS2(MCS):rtTA2_pTRETightBI:mCherry- OPTN_EGFP-LC3b was then cut with SalI and ZraI, and most of it moved into the SmaI and XhoI sites of a derivative of the previous pCS2 (1kb Isl2b) vector containing both 1kb of the RGC-specific Isl2b promoter as well a Flip-recombinase excisable Kanamycin antibiotic selection cassette (pCS2(1kb Isl2b):GFP3-Frt-Kan-Frt; Watson et al., 2012), into which a SmaI restriction site had been previously added immediately downstream of HindIII, so as to generate pCS2(1kb Isl2b):rtTA2_pTRETightBI:mCherry-OPTN_EGFP- LC3b_FKF. All the OPTN mutations (E50K, M98K, F178A, D474N, E478G, and H486R) were then introduced into this construct by QuikChange® Site-Directed Mutagenesis Kit (Stratagene) using the corresponding primers (Primer List below, mutagenized bases are shown in uppercase) and verified by Sanger sequencing. The final step of placing all these under control of the full 20kb of the zebrafish Isl2b promoter, which drives RGC-specific expression in both zebrafish (Pittman et al., 2008) and *X. laevis* (Watson et al., 2012), was accomplished by recombineering (Chan et al., 2007), carried out with minor modifications (Watson et al., 2012).

pCS2(Isl2b:Tom20-mCherry-Apex2_msSOD2UTR): First, to improve the mitochondria targeting efficiency of the transmitophagy reporter, pCS2(1kb Isl2b):Mito-EGFP-mCherry (Davis et al., 2014), the SV40pA was replaced by the 3’UTR of msSOD2 (Kaltimbacher et al., 2006) amplified from mouse brain cDNA (Clontech 637301) using primers AA35 and AA36 and inserted into the XbaI and NotI sites. Then, the mitochondria targeting sequence from Cox8 in this construct was replaced first by Sncg amplified from the same mouse brain cDNA with AA41 and AA42, using HindIII and BamHI, and from there replaced again by the mouse Tom20 sequences amplified from the same mouse brain cDNA with AA47 and AA48, using XmaI and BamHI. To make the Tom20-linker-mCherry-linker-FlagApex2-msSOD2UTR, all relevant fragments were combined in a 4- way Gibson Assembly using the following primers: CD410 and CD621 (Fragment 1), CD620 and CD622 (Fragment 2), CD623 and CD627 (Fragment 3), and CD626 and CD411 (Fragment 4). This construct was originally made to contain within the first fragment a *Xenopus* Rhodopsin promoter (*xop*; (Zhang et al., 2008)), but then the entire cDNA along with the polyadenylation sequence was moved with HindII and NotI into a construct with 1kb of the zebrafish derived Isl2b promoter and Kanamycin selection cassette, so as to make the final construct, referred to simply as Isl2b:Tom20-mcherry in the text, by recombineering as described above.

pCS2(xtFapb7):mTagBFP2-Ras: BFP from pBAD-mTagBFP2 (Addgene plasmid # 34632 ; http://n2t.net/addgene:34632 ; RRID:Addgene_34632) (Subach et al., 2011), was amplified using FK99 and FK98 and cloned into HindIII and BglII sites of a pCS2 vector. The oligonucleotides encoding BglII-Ras farnesylation sequence (AAGCTGAACCCTCCTGATGAGAGTGGCCCCGGCTGCATGAGCTGCAAGTGTGTGCTCTCCTGA) -XbaI was inserted into BglII and XbaI sites of the vector by using complementary oligonucleotides. Then the full cDNA of mTagBFP2-Ras and SV40 polyadenylation sequence was placed in between HindIII and NotI sites of pCS2(xtFabp7) vector.

pCS2(xtFapb7):xlAqp4-GFP: a Aqp4 cDNA was amplified from tadpole optic nerve mRNA using primers NMD 18 and NMD 20 and cloned into HindIII and NheI sites of a pCS2(xtFabp7) construct (Mills et al., 2015) with a unique NheI site in frame upstream of GFP. This DNA was then used as template for PCR reactions using NMD 29 and CD411, and NMD 30 and CD 410, in order to mutagenize the first stop codon of Aqp4, into a tryptophan, through a Gibson ligations of the two amplicons.

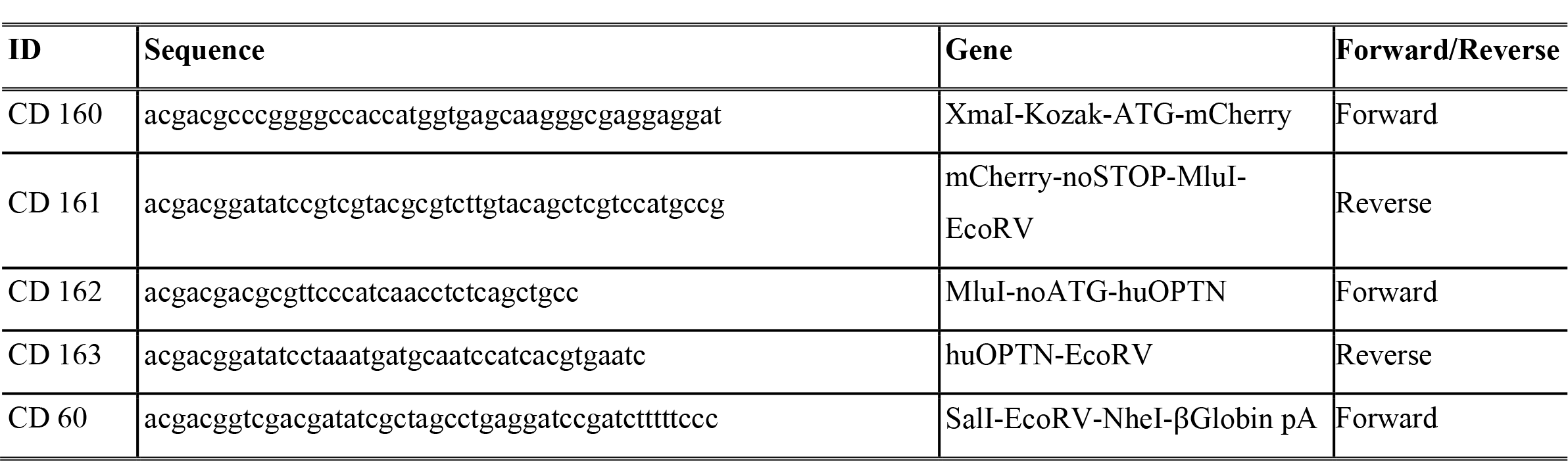
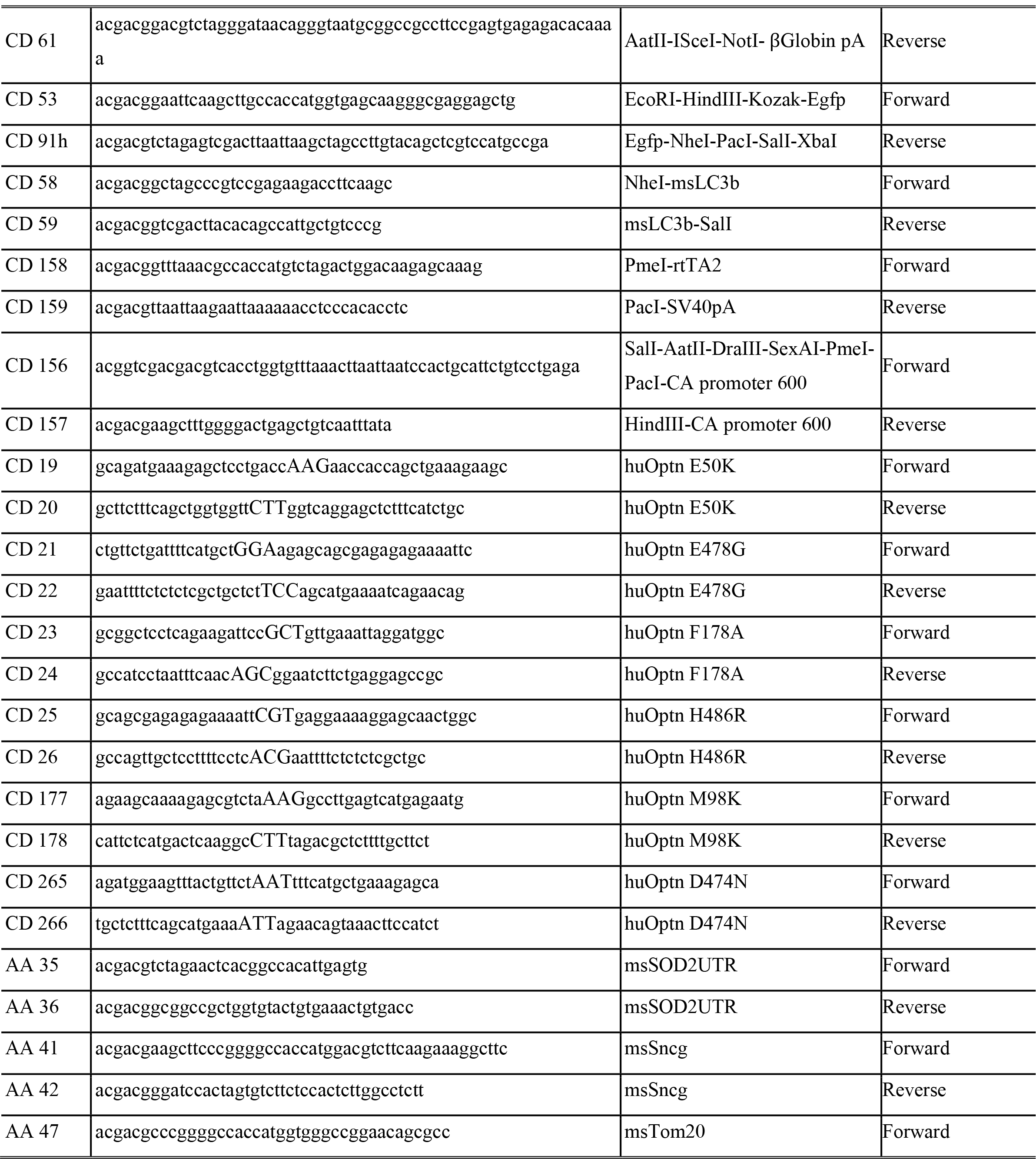
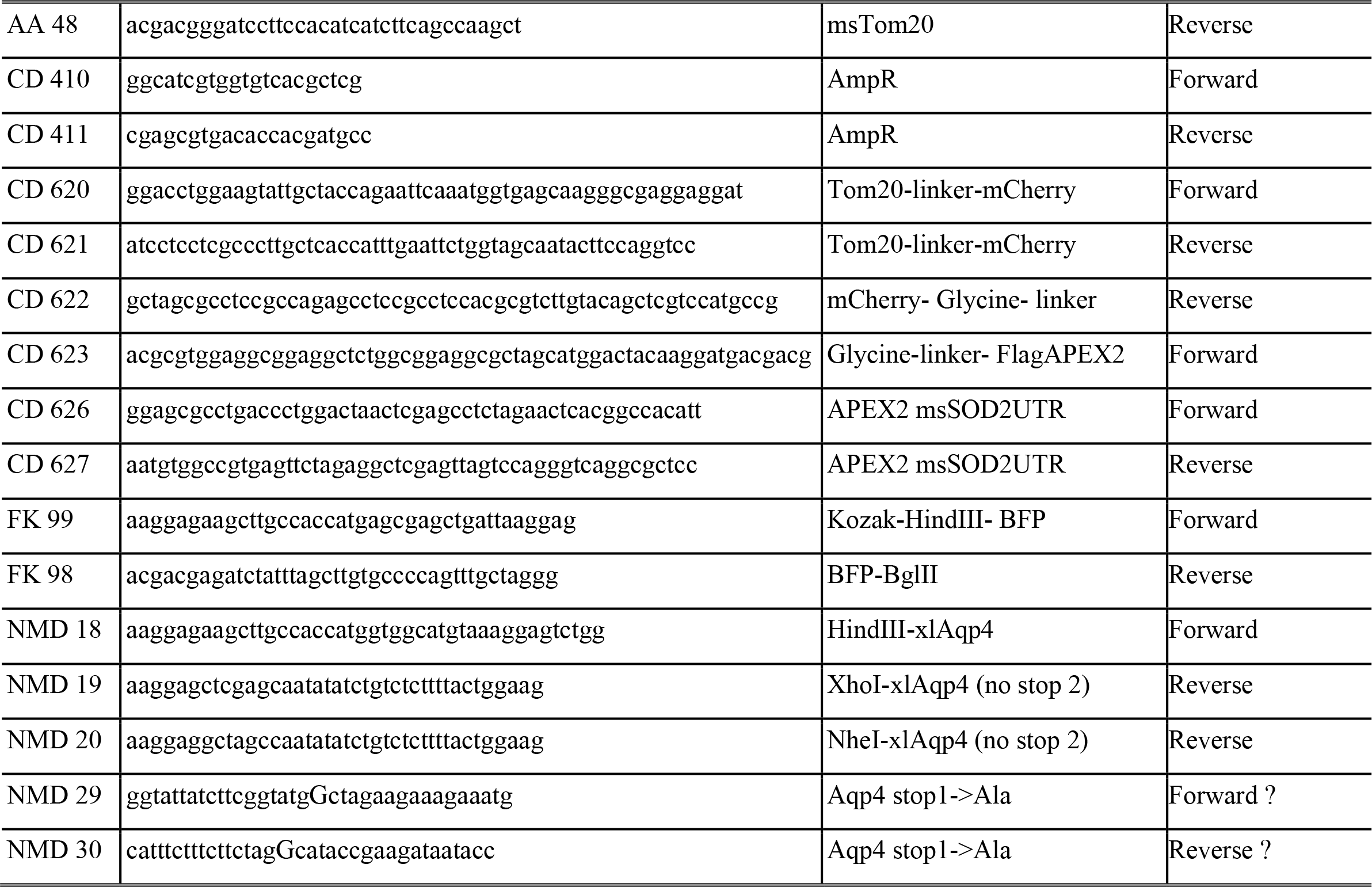

### Transgenesis

Transgenic *X. laevis* were generated by REMI transgenesis (Amaya and Kroll, 1999) using optimizations for larger DNA constructs that have been described previously (Mills et al., 2015). In all cases, DNAs were linearized overnight with NotI in the injection buffer, followed by 65 °C 20-minute inactivation of the enzyme. Embryos were kept in 0.1X Marc’s Modified Ringer’s (MMR) with 50 μg/mL of gentamicin in sterile glass petri-dishes for the first 2 days, the first day at 16 °C and the second at room temperature, and then subsequently kept in 0.1X MMR in glass bowls at room temperature and under a 12/12 light/dark cycle.

### Generation of optic nerves with sparsely labeled axons

Approximately 10-50 eye anlage cells were surgically transferred from *X. laevis* homozygous or heterozygous for Isl2b:mCherry-OPTN(Wt/E50K/M98K)_EGFP-LC3b into either wildtype or *Tg(xtFabp7:mTagBFP2-Ras)* hosts at a developmental stage prior to retinal cell differentiation (at between NF stage 24 and 32). Surgeries were carried under Zeiss Stemi 2000 stereomicroscopes using hand-held curved tungsten micropins on animals anesthetized with 0.2 g/L of Tricaine Methanesulfonate (MS-222) in filter sterilized 0.1X MMR 50 μg/mL of gentamicin with additional buffering provided by 2 mM HEPES, pH 7.4-7.6 (sterile 0.1X MMR), in 35-mm petri dishes. Animals were immobilized during surgeries in disposable clay-mold (Permoplast, non-toxic) inserts with adjacent grooves created with a pipette tip and then altered with forceps to loosely anchor donors and hosts in parallel near each other. At approximately 10 minutes after transplantation, the animals were released from their clay-mold enclosures and donors and hosts returned together to the same dish containing fresh sterile-filtered 0.1X MMR 2mM HEPES with 50 μg/mL of gentamicin. Successful transplants, which ranged from 30-60%, were defined as those with donors confirmed as transgenic (since operations are done prior to onset of transgene expression), and whose hosts had eyes of normal size and only a few RGC axons labeled, as primarily assessed under a Leica MZ10F fluorescence-equipped stereomicroscope and then verified through confocal live-imaging (see details below).

### Intravitreal injection of Mitotracker

Mitotracker Deep Red is a far-red fluorescent dye that chemically stains mitochondria in live cells (Audano et al., 2021; Weiss-Sadan et al., 2019). Fresh 200 µM of MitoTracker® Deep Red FM (Cell Signaling Technology, #8778) dye solution was prepared by diluting a 5 mM of stock solution prepared in DMSO with filtered 0.5X MMR on the day of injection. The Mitotracker solution was front loaded into a borosilicate glass capillary (World Precision Instruments) that had been pulled and then broken to 1-2 µm tip diameter using fine forceps, and then injected intravitreally into left eyes of anesthetized animals at around NF stage 48 (Nieuwkoop and Faber, 1994), using a pressure injector (Narishige IM-300 Microinjector); successful injections were verified by observing an acute but mild swelling of the eye at the time of injection. After injection, the animals were returned to a new dish containing 0.1X MMR at room temperature and imaged 3.5 hours after injection for S1A and S1B and 15-18 hours after injection for all the other experiments.

### Optic nerve live imaging

In the case of doxycycline-inducible transgenes, *X. laevis* tadpoles were placed in 0.1X MMR with 50 μg/mL of doxycycline for 2-3 days before imaging. At around NF stage 48, *X. laevis* tadpoles were anesthetized in 0.1X MMR with 0.2 g/L of MS-222 and mounted into custom Sylgard 184 silicone elastomer (World Precision Instruments) molds. Molds were created using glutaraldehyde fixed tadpoles at the same stages slightly angled using insect pin inserts so as to make left optic nerves parallel to the imaging plane, and then modified with surgical blades to provide an opening before the mouth to facilitate breathing. Tadpoles immobilized in the molds were weighed down by an 18 mm circular coverslip and imaged from beneath in a 35 mm dish modified with a 22 mm square glass coverslip as bottom. Optic nerves of the animals were imaged by spinning disk confocal microscopy (Dragonfly 503 multimodal imaging system, Andor Technology, Belfast, UK) with a 40×/1.10 (magnification/numerical aperture) HC PL APO water immersion objective, using a Leica DMi8 inverted microscope (Leica, Wetzlar, Germany), an iXon Ultra 888 EMCCD camera (Andor Technology, Belfast, UK), Fusion Software (Andor Technology, Belfast, UK) and laser lines of 100 mW 405 nm, 50 mW 488 nm, 50 mW 561 nm, and 100 mW 643 nm; first a time-series was obtained at a single focal plane imaged for 1 min at 1 Hz, and the entire thickness of the nerve at that same location was z-scanned at 1 µm steps. Movies at 7 fps and images were created using FIJI ImageJ.

For some sparsely labeled axons, 5-10 µm thick regions of the optic nerve were subjected to repetitive z-scan live-imaging either at 30 seconds intervals for 5 minutes or at 2 minutes intervals for 10 minutes. Acquired images were saved as separate image sequence files using FIJI ImageJ and imported into IPLab software (Scanalytics). Regions of interests sampled at different timepoints were then registered and quantified using a custom script written in IPLab. All the relevant movies (3 fps for M98K and 1fps for E50K movies) and images were created using FIJI ImageJ.

### Kymograph analyses

Axonal movements of mitochondria, OPTN and LC3b: acquired time-series images were saved as separate image sequence files in FIJI ImageJ after being processed by HyperStackReg or StackReg Plugin to eliminate object drift. The aligned image sequence files were restacked using IPLab software, followed by either manual (for the sparsely-labeled axons and curved axons) or semi-automatic tracing using IPLab custom scripts so as to visualize and measure fluorescence intensity changes over time in the form of kymographs. For the semi-automatic tracings, the outer contour of the optic nerve was traced, and the approximately 30-µm diameter nerve was automatically divided into 0.9 to 1.8 µm parallel swaths, which contained small stretches of axons running in parallel; using such sampling distance and thickness, the same moving objects within axons were not multiply counted, but some of the larger immotile objects might have been multiply counted. Since there was high concordance between ratios of moving and static objects and the velocities of the moving objects when comparing whole nerve semi-automatic kymographs and the more conventional single axon tracing used in the sparse labeling experiments, any error introduced by the novel semi-automatic tracing of axons was deemed minimal. Axonal movement was determined by counting the percentage of moving and stopped objects and measuring the speed of moving over the 1-min imaging period. Objects were classified as moving if their velocity was 0.1 μm/s or faster. Ratios and average speeds were calculated in Excel.

Colocalization Measures: the same aligned and stacked images from the analysis of axonal movements were reprocessed and analyzed by a separate colocalization script but by the same logic through IPLab as described previously, except displaying multiple-channel fluorescence images along the merged color images, and sequentially tracing objects labeled by multiple fluorophores first, followed by those labeled by individual fluorophores. Ratios and average speeds were calculated in Excel.

### Nerve 3D reconstruction and quantification in Imaris

Z-scan confocal images of optic nerves expressing mCherry-OPTN and EGFP-LC3b were imported into Imaris software (Bitplane). Because LC3b is localized not only in autophagosomal membrane, but also is observed more diffusely within cytosol, including axoplasm (Fu et al., 2014; Wong and Holzbaur, 2014b; Tammineni et al., 2017; Stavoe et al., 2019; Evans and Holzbaur, 2020; Kuijpers et al., 2021; Boecker et al., 2021), this EGFP-LC3b signal was used to create a mask to define the boundary of the RGC axons and to separately measure the mCherry-OPTN or far-red mitochondria signal that co-localized or not with the axons. The Imaris mask toolset was used to produce channels for OPTN, mitochondria or OPTN-mitochondria either colocalizing or not with LC3b. The % outside axon values were calculated in Excel by comparing the volumes of objects outside of the LC3b mask relative to the total volumes.

### Correlated Light SBEM

After optic nerve live-imaging, tadpoles were fixed overnight with 4% paraformaldehyde, re-imaged for confocal imaging and positioning, then fixed overnight with 2% EM grade glutaraldehyde (18426, Ted Pella Incorporated) in 0.15M sodium cacodylate buffer (SCB) containing 2 mM CaCl2. The tadpoles were then prepared for serial block-face imaging (SBEM) as described previously (Deerinck, et al., 2010). Briefly, tadpoles were washed with SCB then incubated in 2%OsO4 + 1.5% potassium ferrocyanide in SCB containing 2mM CaCl2 for 1 hour. Tadpoles were washed with double distilled water (ddH2O) and incubated in .05% thiocarbohydrazide for 30 minutes. Tadpoles were then washed again with ddH2O and stained with 2% aqueous OsO4 for 30 minutes. Tadpoles were again washed in ddH2O and placed in 2% aqueous uranyl acetate overnight at 4C. Tadpoles were washed again with ddH2O and stained with en bloc lead aspartate for 30 minutes at 60C. After a last wash in ddH2O, tadpoles were dehydrated on ice in 50%, 70%, 90%, and 100% ethanol solutions for 10 minutes each, followed by two 10 minute incubations in dry acetone. Tadpoles were placed in 50:50 dry acetone/Durcopan resin overnight, followed by 3 changes of 100% Durcopan for ∼12 hours each. Tadpoles were then allowed to harden in Durcopan at 60C for 48 hr.

Tissue blocks were cut from hardened Durcopan, and the whole head of the tadpole was mounted on plastic dummy blocks, trimmed down to only the region surrounding the left optic nerve, eye, and brain. Exact location of optic nerve and the rest the region interest determined by X-Ray microscopy imaging of tissue blocks performed on a Zeiss Xradia Versa 510 X-ray microscope (XRM) instrument (Zeiss X-Ray Microscopy, Pleasanton, CA, USA). XRM tilt series were generally collected at 120 kV and 10W power (75 µA current) and used to create 3D reconstructions of the embedded tissue. SBEM data was collected with a 3View unit (Gatan, Inc., Pleasanton, CA, USA) installed on a Gemini field emission SEM (Carl Zeiss Microscopy, Jena, Germany). Volume was collected at 2.5 kV accelerating voltage, with a raster size of 20k × 17k and pixel dwell time of 1.0 µsec. The pixel size was 5.0 nm and section thickness was 60 nm. SBEM volumes were analyzed and annotated in 3Dmod (3dmod Version 4.11.24, Copyright (C) 1994-2021 by the Regents of the University of Colorado).

### Statistics

Statistical analyses involved the comparison of means using unpaired, two-tailed Student’s t-tests or two-way ANOVAs following Tukey’s post-hoc test for multiple comparisons using GraphPad Prism software. All the bar graphs represent the means ± SEM, where values were first averaged per animal and N is the number of animals. In the stacked bar graphs (except Fig. 2G, S1E, S2H and S2I) showing percentage of each movement, all statistical comparisons were performed using all the axonal movements including stationary, orthograde and retrograde, although only the statistically significant comparisons between the stationary groups (against Wt OPTN or control group) are shown. In all the velocity graphs (except Fig. S1F), all statistical comparisons were performed throughout all the transgenic lines and movements (both orthograde and retrograde movements), and only the statistically significant comparisons against the Wt OPTN or control group are shown.

All animal experiments were carried out in accordance with procedures approved by the Institutional Animal Care and Use Committee of University of California, Davis.

## Supporting information

Mov. 1

Mov. 6

Mov. 5

Mov. 4

Mov. 3

Mov. 2

## Acknowledgments

This work was supported by the National Eye Institute R01EY026471 and R01EY029087) (N.M- A). Correlated light, XRM and Volume EM analysis was performed at the National Center for Microscopy and Imaging Research, with support from NIH grants U24 NS120055, 1S10OD021784 and National Science Foundation - NSF2014862-UTA20- 000890 (M.H.E.). Deposition and management of Volume EM data within the Cell Image Library was further supported by NIH grant R01 GM82949 (M.H.E.). The authors thank Elizabeth A Mills and Ferdinand Kaya for their initial work on generating the original Isl2b- and Fabp7- promoter constructs. Image analyses were performed through the use of UC Davis Health Sciences Advanced Imaging Facility supported by the NEI UC Davis Core grant (P30-EY012576).

## Author contributions

Conceptualization, Y.J., C.-h.O.D., N.M.-A.; Methodology, Y.J., C.-h.O.D., N.M.-A., A.M., M.E.; Software, N.M.-A.; Validation, Y.J., N.M.-A.; Formal Analysis, Y.J., N.M.-A., A.M., M.E.; Investigation, Y.J., C.-h.O.D., N.M.-A., V.D., A.M., M.E.; Resources, N.M.-A., M.E.; Data Curation, Y.J., C.-h.O.D., N.M.-A., A.M., K-Y. K.; Writing-Original Draft, Y.J., N.M.-A.; Writing-Review & Editing, Y.J., C.-h.O.D., N.M.-A., A.M., M.E.; Visualization, Y.J., N.M.-A.; Supervision, N.M.-A.; Project Administration, N.M.-A.; Funding Acquisition, N.M.-A.

---

**Supplementary Figure 1.**
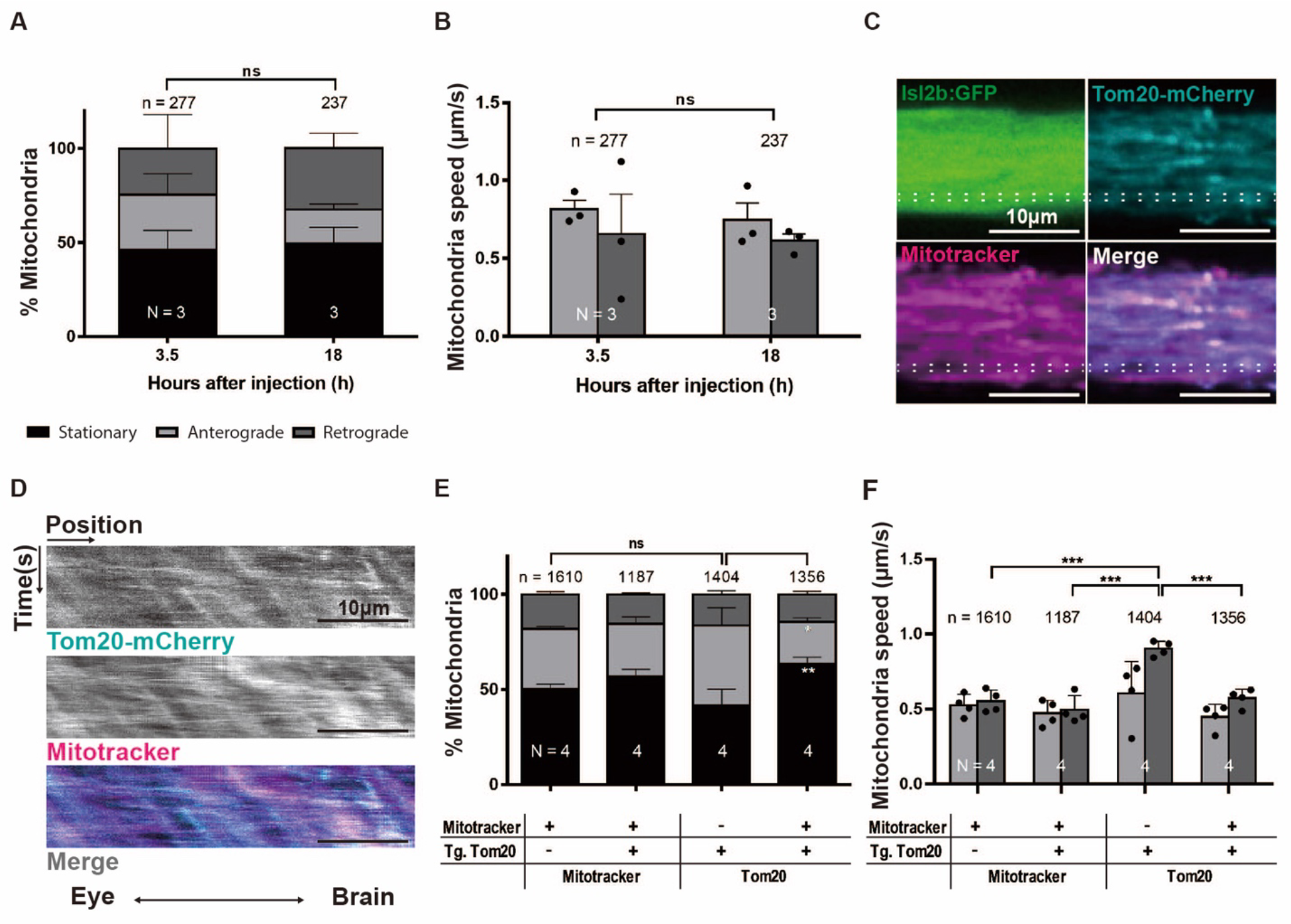
Intravitreal Mitotracker injection results in optic nerve mitochondria labeling that is stable within one day of injection, labels transgenically labeled RGC axonal mitochondria, and has a small but significant effect on the movement behavior of axonal mitochondria. (**A**-**B**) RGC axonal mitochondrial movement metrics are stable within the optic nerve between 3.5 and 18 hours after intravitreal Mitotracker injection. (**A**) Percentage of stationary, and anterogradely and retrogradely moving mitochondria and (**B**) axonal mitochondrial average speed in each direction (anterograde and retrograde). Mean ± SEM; n = 237-277 mitochondria from 3 animals. (**C**-**D**) A Tom20-mCherry transgene expressed by the RGC-specific *Isl2b* promoter and intravitreal Mitotracker injection both label largely the same mitochondria in the optic nerve; representative images (**C**) and corresponding kymographs (**D**). (**E**-**F**) Quantification of mitochondrial movements in RGC axons based on either Mitotracker or the Tom20-mCherry transgene. From left to right, the measurement based on Mitotracker-labeled objects in Mitotracker injected *Tg(Isl2b:GFP)* animals, Mitotracker-labeled objects in Mitotracker injected *Tg(Isl2b:GFP x Isl2b:Tom20-mCherry)* animals, Tom20-labeled objects in *Tg(Isl2b:GFP x Isl2b:Tom20-mCherry)* animals, Tom20-labeled objects in Mitotracker injected *Tg(Isl2b:GFP x Isl2b:Tom20-mCherry)* animals. Mean ± SEM; n = 1187-1610 mitochondria from 4 animals. Statistical analysis in **A**, **B**, **E** and **F** were performed by two-way ANOVA following Tukey’s post- hoc test for multiple comparisons. Not significant (ns), ** = p<0.01, *** = p<0.001.

**Supplementary Figure 2.**
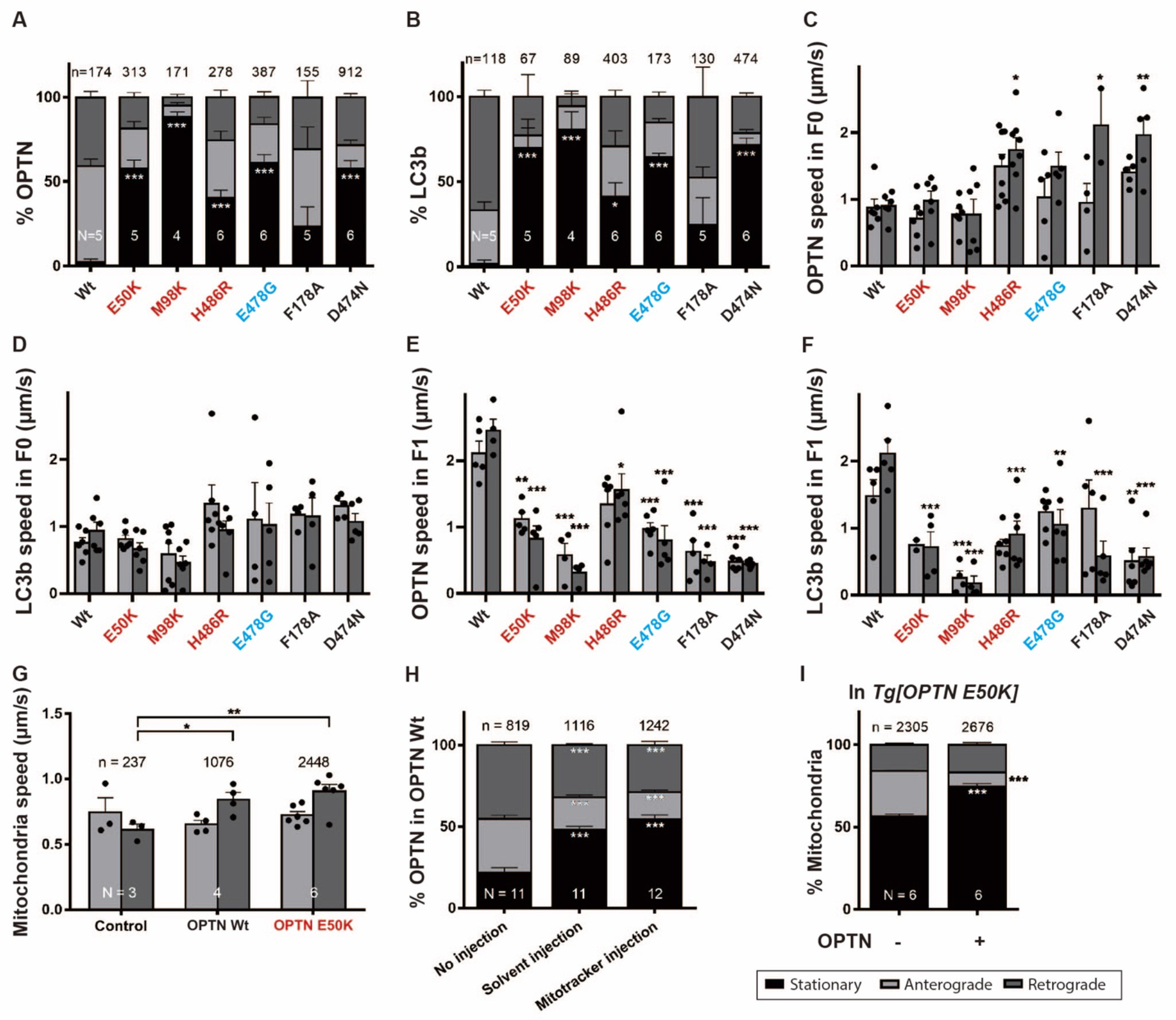
Movement of OPTN and LC3b in F1 animals (not injected intravitreally with Mitotracker) match those observed in F0 animals, including showing no large velocity changes in either orthograde or retrograde movement. Percentage of each movement (stationary, anterograde and retrograde) of OPTN (**A**) and LC3b (**B**) in F1 transgenic animals and (**C**-**F**) average speed of axonal OPTN and LC3b in each direction (anterograde and retrograde) in F0 (**C**- **D**) and F1 (**E**-**F**) transgenic animals. Mean ± SEM; n = 155-912 OPTN from 4-6 animals. n = 67- 474 LC3b from 4-6 animals. (**G**) Average anterograde and retrograde speed in control (non- transgenic, only Mitotracker-injected animals), and Wt OPTN and E50K OPTN expressing RGC axons. Mean ± SEM; n = 237-2448 mitochondria from 3-6 animals. (**H**) Fraction of stopped and moving OPTN in animals expressing Wt OPTN in the presence and absence of intravitreal injections (0.5X MMR solvent or Mitotracker injections). The Mitotracker dye does not affect an increase in the percentage of stationary objects; rather, it is the intravitreal injection itself that causes the augmentation. Mean ± SEM; n = 819-1242 OPTN objects measured in 11-12 animals. (**I**) Quantification of mito-OPTN colocalizations in Mitotracker-injected E50K OPTN animals. Percentage of each movement (stationary, anterograde and retrograde) of mitochondria co- localizing with (mito-OPTN) or not co-localizing with (mito-ONLY) OPTN in E50K OPTN expressing animals. Mean ± SEM; n = 2305 mito-ONLY, n = 2676 mito-OPTN colocalizations from 6 E50K OPTN animals. Statistical analysis in **A-I** were performed by two-way ANOVA following Tukey’s post-hoc test for multiple comparisons * = p<0.05, ** = p<0.01, *** = p<0.001.

**Supplementary Figure 3.**
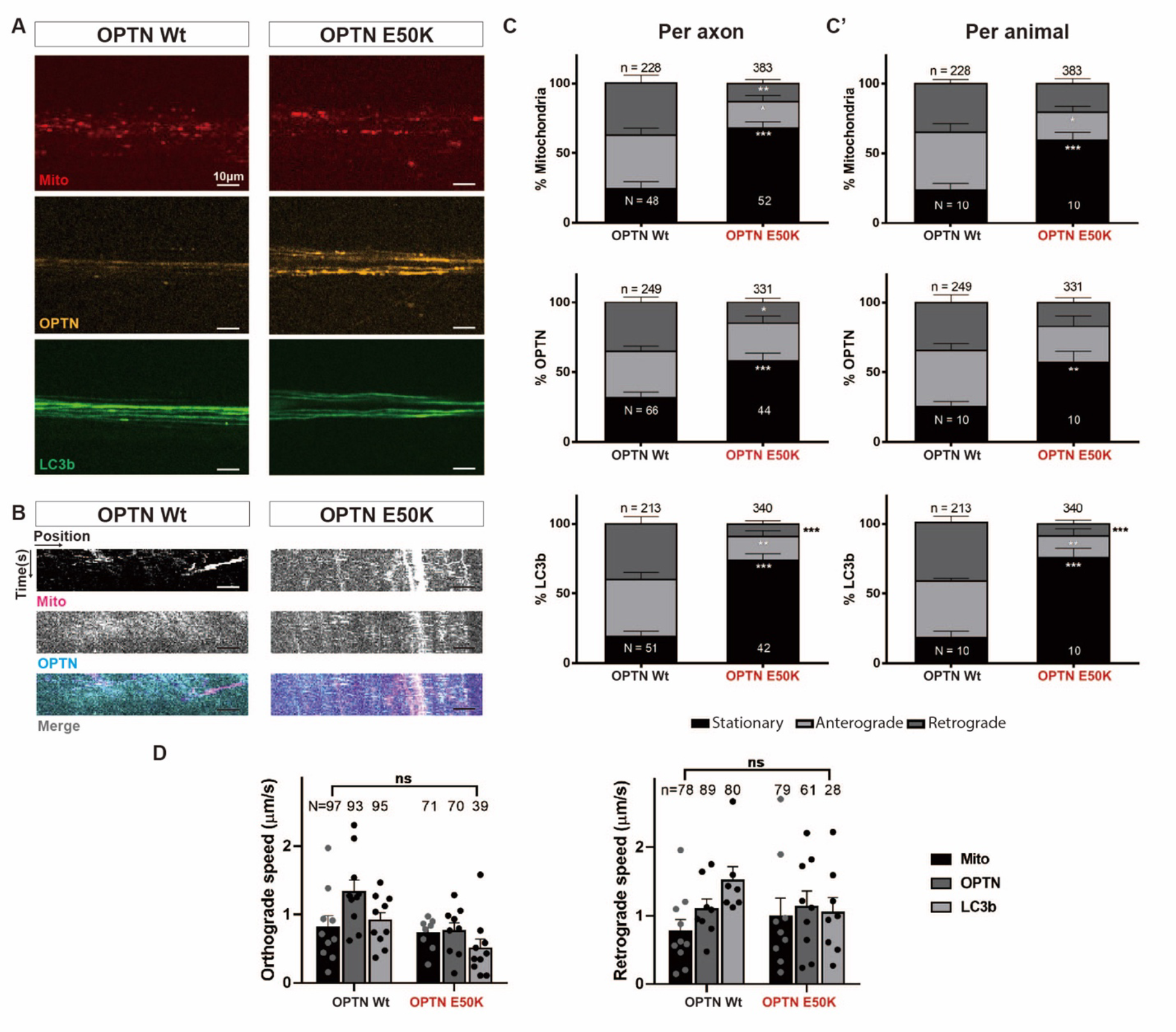
Movements of axonal mitochondria, OPTN and LC3b within the sparsely labeled Mitotracker labeled *Tg(Isl2b:mCherry-OPTN(Wt or E50K)_ EGFP-LC3b)* axons of RGCs transplanted into non-transgenic *X. laevis*. (**A**-**B**) Representative confocal images (**A**) and corresponding kymographs (**B**) of axonal mitochondria and OPTN within the sparsely-labeled axons. Scale bar, 10 µm. (**C-C’**) Relative to Wt OPTN, E50K OPTN increases the percentage of stationary mitochondria, OPTN and LC3b measured in sparsely labeled axons (analyzed per axon (**C**) and per animal (**C’**)). Mean ± SEM; n = 228-383 mitochondria in 48-52 axons from 10 animals. n = 249-331 OPTN in 44-66 axons from 10 animals. n = 213-340 LC3b in 42-51 axons from 10 animals. (**D**) Average speed of axonal mitochondria, OPTN and LC3b in each direction (anterograde and retrograde) in Wt and E50K OPTN transgenic animals. Mean ± SEM; n = 71-97 mitochondria in 48-52 axons from 10 animals. n = 61-93 OPTN in 44-66 axons from 10 animals. n = 28-95 LC3b in 42-51 axons from 10 animals. Statistical analysis in **C-D** were performed by two-way ANOVA following Tukey’s post-hoc test for multiple comparisons. Not significant (ns); * = p<0.05, ** = p<0.01, *** = p<0.001.

